# A genome-scale metabolic model of *Drosophila melanogaster* for integrative analysis of brain diseases

**DOI:** 10.1101/2022.08.22.504778

**Authors:** Müberra Fatma Cesur, Kiran Raosaheb Patil, Tunahan Çakır

**Author notes:** Correspondence: Tunahan Çakır.

## Abstract

High conservation of the disease-associated genes between fly and human facilitates the common use of *Drosophila melanogaster* to study metabolic disorders under controlled laboratory conditions. However, metabolic modeling studies are highly limited for this organism. We here report a comprehensively curated genome-scale metabolic network model of *Drosophila* using an orthology-based approach. The gene coverage and metabolic information of the orthology-based draft model were expanded via *Drosophila*-specific KEGG and MetaCyc databases, with several curation steps to avoid metabolic redundancy and stoichiometric inconsistency. Further, we performed literature-based curations to improve gene-reaction associations, subcellular metabolite locations, and updated various metabolic pathways including cholesterol metabolism. The performance of the resulting *Drosophila* model, termed iDrosophila1 (8,230 reactions, 6,990 metabolites, and 2,388 genes), was assessed using flux balance analysis in comparison with the other currently available fly models leading to superior or comparable results. We also evaluated transcriptome-based prediction capacity of the iDrosophila1, where differential metabolic pathways during Parkinson’s disease could be successfully elucidated. Overall, iDrosophila1 is promising to investigate systems-level metabolic alterations in response to genetic and environmental perturbations.

## 1 Introduction

*Drosophila melanogaster* is a well-known model organism with highly tractable genetics in order to get insight into human metabolism. It contains the counterparts of various essential human systems such as central nervous system, gastrointestinal system, kidney (Malpighian tubule in fly), adipose tissue (fly fat body), and liver (fly oenocytes) [1]. In addition to this conservation, shorter lifespan (transition from embryo to adulthood within 10-14 days), rapid generation time, the large numbers of progeny, a well-defined genome, substantially less genetic redundancy, and the availability of advanced genetic tools for this organism encourage researchers to investigate many aspects of human diseases via *Drosophila* [2, 3]. The highly conserved disease pathways of human have been extensively analyzed based on the *Drosophila* genes, which share 75-77% homology with disease-associated human genes [3, 4].

Genome-scale metabolic network (GMN) models facilitate the mathematical representation of biological knowledge about cellular metabolism through the inclusion of all known chemical reactions, metabolites, and genes for an organism. The genes are linked to the associated reactions via conditional statements in Boolean logic, which are known as gene-protein-reaction (GPR) rules [5]. Metabolic models are promising to address many scientific questions and introduce novel avenues for the identification of potential drug targets, detection of putative biomarkers, *in silico* metabolic engineering, pan-reactome analyses, understanding of metabolic disorders, and modeling of host-pathogen interactions [5–7]. Therefore, a wide variety of GMN models have been so far developed for prokaryotic and eukaryotic model organisms [5]. In spite of impressive experimental efforts and developing technologies, there is only one recent metabolic model representing comprehensive *D. melanogaster* metabolism [8]. This model, called Fruitfly1 (12,056 reactions, 8,132 metabolites, and 2,049 genes), was derived from generic human GMN (Human1) [9] based on gene orthology information from the Alliance of Genome Resources [10]. Another comprehensive *Drosophila* model (BMID000000141998) was developed in 2013 without any manual curations [11]. It consists of 6,198 reactions, 2,873 metabolites, and 4,020 genes. Despite the high gene coverage of this model, it allows the synthesis of all biomass components even without any carbon source consumption by the model [11, 12]. In addition, two curated small-scale metabolic models (flight muscle and larvae models) are available for this organism. They only represent the core metabolism of *Drosophila* [12, 13]. The flight muscle model (194 reactions, 188 metabolites, and 167 genes) was developed in 2007 to elucidate cellular adaptation mechanisms against hypoxia and it was revised in 2008 [13, 14]. More recently, the second metabolic model called FlySilico (363 reactions, 293 metabolites, and 261 genes) was built to simulate larval development [12]. These models are not suitable to investigate the complex metabolism of fruit fly.

Comprehensive metabolic network models can be developed considering genetic similarity between organisms. This semi-automated reconstruction approach starts with the generation of a draft model based on a template model. The template model has a high genetic similarity with the organism of interest. The genes of this reference network are replaced by their orthologous counterparts in the target organism. Thus, existing information about gene associations is transferred to reconstruct new GMN models [15]. To date, orthology-based GMN models have been developed for several organisms thanks to the high degree of genomic homology between human and these model organisms [8, 15–17]. In the current study, we developed a GMN model of *D. melanogaster* using a curated version of HMR2, a human GMN model [18], as the template. To do so, we first developed a draft model through the orthology-based mapping of *Drosophila* genes. Then, the draft *Drosophila* model was expanded based on metabolic information in KEGG and MetaCyc databases. Manual curation steps were performed for each model component in several steps of the reconstruction process. Thus, we reconstructed a comprehensively curated *D. melanogaster* model called iDrosophila1. This GMN model was analyzed in terms of its phenotypic predictive capacity and gene essentiality prediction. Furthermore, iDrosophila1 was shown to represent metabolic alterations corresponding to Parkinson’s disease (PD). Taken together, we believe that iDrosophila1 can enable extensively characterization of fly metabolism as well as investigation of molecular basis of complex human diseases.

## 2 Material and Methods

### 2.1 Metabolic reconstruction procedure

We used an orthology-based approach to reconstruct a draft *D. melanogaster* model based on a flux-consistent version of human HMR2 model (7,518 reactions, 5,426 metabolites, and 2,479 genes) as the template [18]. We curated the GPR associations of the template model by improving protein complex information based on another human model (iHsa), which includes an extensive manual refinement in the GPR rules of HMR2, especially in terms of defining enzyme complexes [17]. The orthology-based inference of *Drosophila* model was achieved through the replacement of genes in the GPR rules of template model with *Drosophila* orthologs. To do so, ‘human orthologs’ list of *D. melanogaster* was retrieved from the FlyBase database (last accessed: 02/02/2021) [19]. The list includes evidence scores called DIOPT scores in addition to information about *Drosophila* genes (FlyBase gene IDs), their human orthologs, and associated disease phenotypes. DIOPT score refers to the number of tools supporting a given orthology prediction [20]. It was used in orthology-based gene mapping as a scoring approach (3 to 15) to increase the levels of confidence (Supplementary Table 1). In this approach, two criteria were considered: (1) If a human gene has multiple *Drosophila* orthologs with different DIOPT scores, only orthologous gene(s) with maximum DIOPT scores were considered. (2) If a human gene has over nine *Drosophila* orthologs, this gene association was ignored [17] but the related reactions were kept in the model. Lastly, GPR rules of the template model were modified by replacing human genes with their *Drosophila* orthologs at a high confidence level, and the human genes that could not be matched with any *Drosophila* orthologs were also discarded from the model. On the other hand, all non-enzymatic reactions in the template model were included in the draft model.

To enhance gene coverage of the draft model, we generated another metabolic network for *D. melanogaster* using RAVEN toolbox [21] by considering protein homology against KEGG [22] and MetaCyc [23] databases. Based on the study of Wang and colleagues (2018) [21], we first reconstructed a combined KEGG-based metabolic network through merging two different KEGG-based networks: (1) We used only KEGG organism identifier (dme) for the first network and (2) the second network was reconstructed using BLAST algorithm to query the protein sequence of *Drosophila* retrieved from FlyBase database (release number: r6.37) against the latest pre-trained hidden Markov Model (euk100_kegg94). The same parameters (default cut-off: 10^-50^, minScoreRatioG: 0.95, and minScoreRatioKO: 0.7) were used in the reconstruction of both KEGG-based networks, and incomplete information/undefined stoichiometries were excluded. In addition to the combined KEGG-based metabolic network, we generated a MetaCyc-based metabolic network using another automatic reconstruction function in the RAVEN toolbox. In the reconstruction process, default parameters (bit-score ≥ 100 and positives ≥ 45%) were selected and unbalanced/undetermined reactions were excluded from the network.

KEGG-and MetaCyc-based metabolic networks were subsequently merged (hereafter referred to as KEGG-MetaCyc network). The genes that were not shared with the orthology-based draft model were identified. These KEGG-MetaCyc-specific genes were functionally characterized through the identification of significantly enriched KEGG pathways using g:Profiler web server [24] with a false discovery rate (FDR) at 0.05 level (Supplementary Table 2). The reactions associated with the KEGG-MetaCyc-specific genes were selected to be included in the draft model. Due to the lack of compartmentalization in the KEGG-MetaCyc network, subcellular protein localizations reported by several sources were used to assign at least one compartment for each KEGG-MetaCyc-specific gene (Supplementary Table 3). The sources utilized in this study include COMPARTMENTS [25], FlyBase [19], GLAD [26], QuickGO [27], AmiGO 2 [28], UniProt [29], Reactome [30], CELLO2GO web server [31], and a mass-spectrometry-based study [32]. It is important to note that gene ID consistency is crucial for an accurate gene-compartment mapping. Therefore, FlyBase ID Validator tool [19] was used for the conversion of IDs from the subcellular localization databases and tools to the current versions of the FlyBase gene IDs. Then, a compartment dictionary was generated through mapping the compartment information to the related KEGG-MetaCyc-specific genes based on Gene Ontology (GO) cellular components term. Only compartments found in the draft model were considered while generating this dictionary. After the gene compartmentalization process, the compartment dictionary facilitated the transfer of the compartment information from the genes to the KEGG-MetaCyc reactions based on their GPR associations.

The orthology-based draft *Drosophila* model was merged with the compartmentalized KEGG-MetaCyc-specific model by adding each KEGG-MetaCyc reaction to the draft model using *addReaction* function in COBRA toolbox [33]. The biomass formation equation derived from the template human model was curated, as explained in the next section. We identified biomass components that could not be synthesized by the merged model using COBRA *biomassPrecursorCheck* function. Then, a gap-filling step was employed via *fillGaps* algorithm in the RAVEN toolbox in order to add reactions required for the synthesis of those biomass components. To ensure the production of all biomass components, the lower bound of biomass formation reaction was set to 0.1 while running *fillGaps* algorithm. In addition, we set the parameter, ‘useModelConstraints’ in the *fillGaps* to true. In this process, the template human model was used as the reference to fill in missing metabolic knowledge. First, the reference human model and merged model were set to a chemically defined medium with unlimited uptake rates (1000 mmol g^−1^h^−1^). This growth condition was called as expanded holidic diet (HD) due to the inclusion of several additional vitamin derivatives to the HD medium (see Supplementary Table 4) [12, 34]. Then, new reactions were added to the merged model via gap filling from the human model without gene information (Supplementary Table 5).

After the gap-filling step, cholesterol metabolism was revised to represent the cholesterol auxotrophy of *Drosophila*. In addition, GPR rules were updated based on the information about 556 *Drosophila* protein complexes generated by Guruharsha and colleagues (2011) [35]. Further curations were introduced for the GPR rules based on the FlyBase gene group list and literature [36–48]. We also investigated the presence of leaking energy metabolites in the draft model. In the leak testing, we analyzed the metabolism of charging energy metabolites listed by Fritzemeier and colleagues (e.g., ATP, NADH, NADPH, FADH_2_, GTP, and H^+^) [49]. To this aim, we first constrained all uptake reactions to zero by allowing only secretion in the model. Then, a dissipation reaction was added for each energy metabolite if the related reaction was missing in the model. Each dissipation reaction was defined as the objective function to evaluate whether the corresponding metabolite is leaking or not. Thus, we determined leaking metabolites based on the optimal flux of these reactions. If any dissipation reaction was found to be active under nutrient-deficiency condition, the related energy metabolite was accepted as a leaking compound. To overcome this issue, we performed multiple reaction deletions and identified the reactions causing metabolic leaks. The identified reactions were manually curated. It is worth emphasizing that the reaction ‘HMR_4762’ mediating porphyrin metabolism was also updated to revise the incorrect link between heme and cytochrome-C metabolites. In fact, this reaction is catalyzed by cytochrome-C heme lyase (FBgn0038925) for the conversion of apocytochrome-C and heme to cytochrome-C in a reversible manner. Due to the incorrect conversion of heme to the cytochrome-C in the template human model, we first added apocytochrome-C to HMR_4762 in the *Drosophila* model for the proper conversion. Then, the missing reactions associated with the apocytochrome-C metabolism were added from HMR2 model. These modifications have impact on the growth rate due to the presence of cytochrome-C in the ‘cofactor and vitamin’ composition of the biomass equation. One should note that we performed several additional curation steps for all components of the metabolic networks (reactions, metabolites, and genes) considered in the reconstruction steps of the *Drosophila* model. The details of these steps are explained in Supplementary Note. The final *Drosophila* model was called iDrosophila1, and it can be accessed from the GitHub repository (https://github.com/SysBioGTU/iDrosophila).

### 2.2 Curation of biomass formation reaction

The biomass equation of draft *Drosophila* model derived from the reference human model was updated based on FlySilico model [12] and literature. Biomass composition was examined in a comparative manner with FlySilico, and most of the metabolites were determined to be common (amino acids, glycogen, triglyceride, and cholesterol). Missing growth-associated ATP maintenance reaction was included in the model with the coefficient specified in FlySilico. Furthermore, cardiolipin was removed from the biomass reaction because it was reported to be non-essential in *D. melanogaster* [50]. Vitamin D was also removed from the biomass reaction since it is synthesized from 7-dehydrocholesterol, a cholesterol precursor, and, being known as a cholesterol auxotroph, the fruit fly does not have associated enzymes for vitamin D synthesis (See Section 3.2 for details).

### 2.3 Validation of growth phenotypes

iDrosophila1 model was checked for the nonzero production of all biomass precursors. Once we ensured nonzero growth rate by the network, we analyzed cholesterol auxotrophy and the non-essentiality of aspartate amino acid under expanded HD condition (Supplementary Table 4). Here, we allowed flexible intake of the 47 dietary compounds by limiting the maximum sucrose uptake rate to ∼ 2.212 mmol g^−1^h^−1^ [12]. Maximum vitamin uptake rates were set to 1/100 of the sucrose uptake rate due to the low vitamin consumption tendency of organisms while the use of remaining HD substances (except for salts and water) were set to 1/10 of the sucrose uptake rate. We constrained maximum oxygen uptake rate to 24 mmol g^−1^h^−1^ by estimating the oxygen level required to consume all sucrose through aerobic respiration. Under this condition, we investigated the effect of increasing cholesterol and aspartate levels on growth rate as explained in the study of Schönborn and colleagues [12]. To predict the growth rates, maximum biomass production was defined as the objective function and flux balance analysis (FBA) approach [51] was used to identify intracellular flux distributions.

### 2.4 Validation of essential gene predictions

To discover vital genes in iDrosophila1 model, gene essentiality analysis was performed under the expanded HD condition by allowing infinite intake of all diet compounds. Each gene was deleted by suppressing the corresponding reactions, and FBA was performed under the objective of growth maximization. This step was achieved using *singleGeneDeletion* function in COBRA toolbox [52]. The effect of each single-gene knockout on the biomass formation was assessed based on the specified cut-off value (1% of the optimal wild-type growth rate [53]). If the deletion of a gene resulted in a significantly reduced growth rate (i.e., smaller growth rate than the given cut-off), this gene was considered as essential. Thus, we uncovered essential and non-essential gene sets. The genes involved in only inactive (blocked) reactions were subsequently discarded from the list of gene sets due to the absence of their influence to biomass formation. The blocked reactions were identified via flux variability analysis (FVA) [54] under the expanded HD condition. If the sum of absolute minimum and maximum fluxes of a reaction was less than 10^-5^ in FVA, it was accepted as inactive. After determining essential and non-essential gene sets in the active reactions, we characterized the essential genes (Supplementary Table 6A) through the identification of enriched biological processes (Supplementary Table 6B) and KEGG pathways (Supplementary Table 6C) using g:Profiler web server for FDR at 0.05 level. In the next step, we assessed the predictive capability of the model based on the experimental gene essentiality dataset, which was stored in Online GEne Essentiality (OGEE) database [55]. In this dataset, we classified conditionally essential genes as essential, as well. Lastly, several metrics (sensitivity, specificity, accuracy, precision, F1 score, and Matthew’s correlation coefficient (MCC)) were calculated to evaluate the model performance based on the OGEE dataset.

### 2.5 iDrosophila1-mediated analysis of differential metabolic pathways in Parkinson’s disease

We further evaluated the capacity of iDrosophila1 model in phenotypic predictions through the investigation of age-and PD-dependent differential pathways in *D. melanogaster*. In this process, we used a microarray (Agilent) dataset from ArrayExpress (accession number: E-MTAB-1406) [56]. It includes the samples from the heads of male flies harboring *pink1^B9^*(*pink1*) and *park^25^* (*parkin*) mutations in different age groups: young flies (3-day-old) and middle-aged flies (21-day-old (*parkin*) and 30-day-old (*pink1*)). For each age group, *pink1* and *parkin* mutant flies have three and six biological replicates, respectively. Limma package [57] for R version 4.1.0 was used to process, normalize, and analyze the data. In the data processing step, we removed the effects of non-specific signals in the dataset using *backgroundCorrect* function after reading the intensity data via *read.maimages* function. Then, *normalizeBetweenArrays* function was used to achieve the consistency between different arrays. Using this normalized log-transformed dataset, differential expression analysis was employed to uncover significant fold change values for each mutant group relative to the corresponding control group (age-matched wild-type fly). To ensure the consistency of the genes with iDrosophila1 model, all gene annotation IDs were converted to the current versions of FlyBase gene IDs via FlyBase ID Validator tool [19]. In this curation step, multiple hits were manually revised. We subsequently used limma-trend function by setting robust = TRUE. Genes with significant alterations in their expression levels (p-value < 0.01) were kept for further analysis. Based on fold change cut-off values, we applied another filtering process for the genes with multiple probe measurements using following criteria: (1) If any probe(s) of a gene have fold changes ≥1.5, the maximum of fold change values of its probes was considered by assuming the upregulation of this gene. (2) If the fold change values of any probe(s) ≤ 0.67 (∼1/1.5) for a gene, the minimum fold change value was assigned by assuming the downregulation of this gene. (3) If all probes of a gene exhibited moderate (0.67 < fold change < 1.5) or ambiguous fold change values (i.e., the presence of both upregulated and downregulated probes), average fold change was assigned to this gene.

In metabolic network analysis step, the maximum uptake rates of all exchange metabolites were set to unconstrained flux (i.e., 1,000 mmol g^−1^h^−1^). Then, the filtered fold change values were mapped to the reactions in iDrosophila1 model through COBRA *mapExpressionToReactions* function. GPR associations were taken into consideration in the mapping process. Minimum fold change value was assigned to the reactions whose corresponding genes are linked with ‘AND’ operator while maximum fold change was used for the genes that are linked with ‘OR’ operator. Differential reaction expression levels were subsequently used to elucidate PD-induced altered metabolism via a recent approach, ΔFBA with default parameters [58]. Note that the flux of non-growth-associated ATP maintenance was assumed to remain unchanged between wild-type and mutant groups. The ΔFBA algorithm computes flux changes (Δv) between two diverse conditions by applying a two-step optimization procedure: it maximizes the consistency and minimizes inconsistency between Δv and differential reaction expressions. Based on the predicted Δv distribution, we identified the altered (upregulated and downregulated) reaction sets with differential fluxes above the given threshold (|Δvi| > 0.1% of the largest flux bound). GPR rules were used to determine the corresponding genes involved in the regulated reactions. These genes were characterized in terms of significantly enriched KEGG pathways (FDR < 0.05), and diseases (p-value < 0.01) using g:Profiler [24] and FlyEnrichr [59] web servers.

## 3 Results and Discussion

Here, we developed a comprehensive genome-scale metabolic network model for *D. melanogaster* using available metabolic information. In the reconstruction process, extensive model curations associated with metabolic redundancy, stoichiometric consistency, gene/compound name standardization, and missing/incomplete components were performed. These curation steps were applied for both the draft metabolic network and the KEGG-MetaCyc metabolic network, and they are explained in Supplementary Note. Thus, we aimed to avoid any inconsistencies and redundancies in the reconstructed networks by revising each metabolic component (reactions, metabolites, and genes). Additional curations were also applied, if necessary. Overall model reconstruction process is summarized in Figure 1.

**Figure 1.**
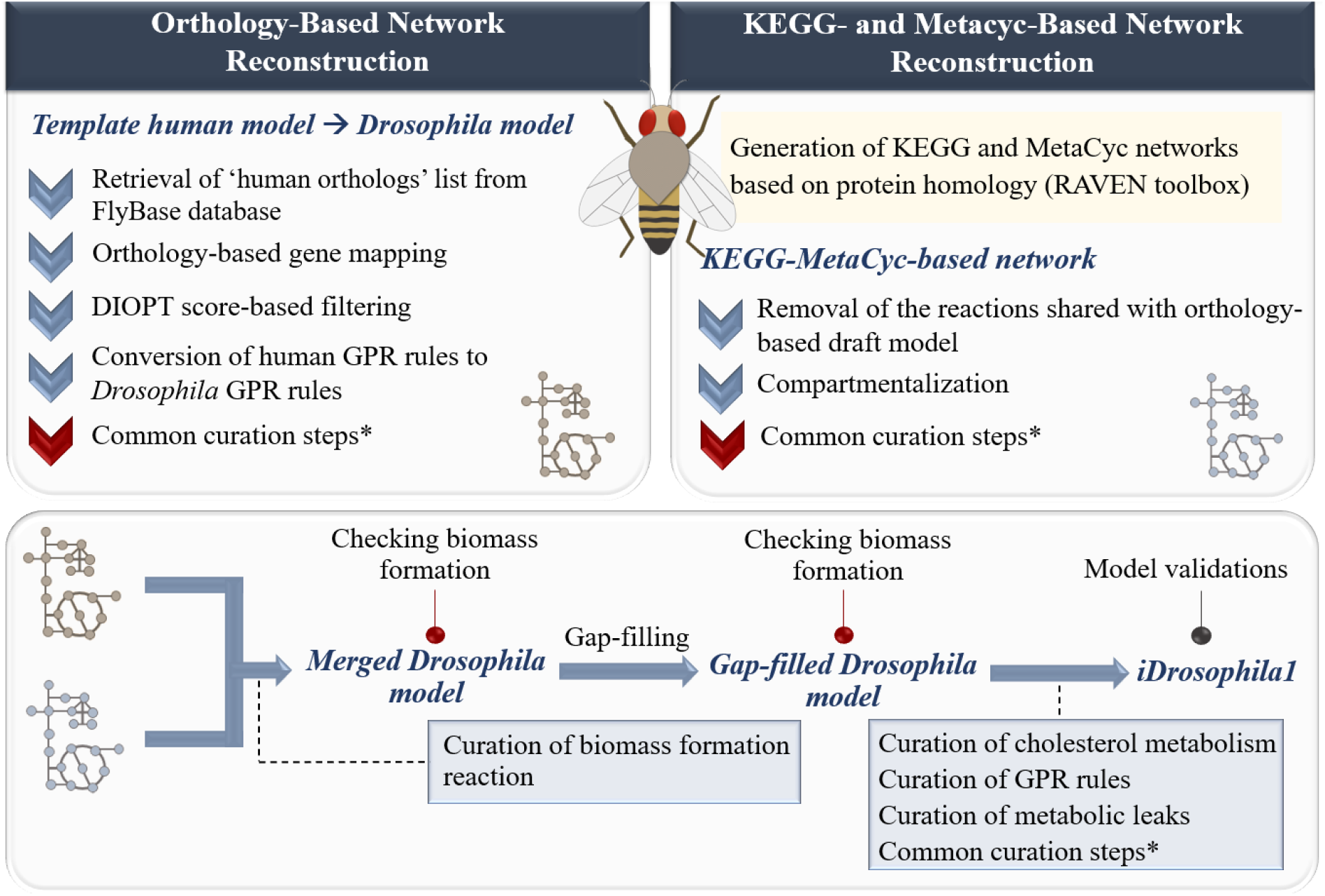
Summarized reconstruction process of the genome-scale metabolic network model for *Drosophila melanogaster*. Commonly applied curation steps are detailed in Supplementary Note.

### 3.1 Draft reconstruction of *Drosophila* metabolic networks

As the starting point, we reconstructed a draft *D. melanogaster* model based on a recent template human model using orthology-based approach. The template model, derived from HMR2 model, contains various improvements such as the addition of mitochondrial intramembrane space, removal of atomically unbalanced reactions, and curation of GPR associations [18]. We further curated the GPR rules based on the information about protein complexes in iHsa model [17]. Thus, over 300 GPR associations were curated. In the reconstruction process of the draft fly model, we used metabolic information in the curated human model. In this process, the GPR rules of the template model were converted to *Drosophila* GPR rules via orthology-based gene mapping at a high confidence level, as explained in Section 2.1. At least one *Drosophila* ortholog was identified for 1,986 out of 2,479 human model genes (Supplementary Table 1). Only one *Drosophila* ortholog was assigned to ∼90% of the human genes, while the remaining genes were matched with multiple orthologs. Presence of the same *Drosophila* orthologs for some human genes led to duplicated *Drosophila* genes in the draft network. These genes were revised to prevent redundancy in the model (Supplementary Note). Gene-associated human model reactions with no *Drosophila* orthologs were not included in the draft model. We also kept the non-enzymatic reactions in the model. This approach enabled the generation of an orthology-based draft *Drosophila* model through the transfer of information about gene associations. The reconstructed draft model includes 6,873 reactions, 4,856 metabolites, and 1,321 genes.

Using metabolic information in KEGG and MetaCyc databases, a KEGG-MetaCyc metabolic network was also generated for *D. melanogaster*. We encountered two major issues associated with this network including (1) a need for the curation of metabolite names and (2) the lack of compartmentalization. As highlighted before, the main aim of considering KEGG-MetaCyc network is to expand gene coverage and available metabolic information in the orthology-based draft model. Therefore, metabolite names in the KEGG-MetaCyc network should be compatible with the draft model for proper merging. To do so, we updated compound names in the KEGG-MetaCyc network by replacing them with the counterparts in the orthology-based draft model (Supplementary Note; also, see Supplementary Figure 1). This step is crucial to reduce a potential metabolic redundancy in the merged model. To avoid metabolic redundancy, we also removed the KEGG-MetaCyc genes that were shared by the draft model, leading to a KEGG-MetaCyc-specific metabolic network. In this step, we determined 1,197 KEGG-MetaCyc-specific genes and 811 common genes. The KEGG-MetaCyc-specific genes were characterized by identifying enriched KEGG pathways. In addition to the fundamental pathways (e.g., carbohydrate, amino acid, fatty acid, and cofactor metabolism), drug and xenobiotics metabolic processes were found among the enriched pathways (Supplementary Table 2). Xenobiotics are any exogenous life-threatening, toxic compounds (e.g., pharmaceuticals, pesticides, and pollutants) to which organisms are exposed [60, 61]. Involvement of the xenobiotics in neurodegenerative disorders like Alzheimer’s disease (AD) and PD was reported [62, 63]. Accordingly, rotenone and paraquat are commonly used to induce a PD-like phenotype (e.g., movement disorders and loss of dopaminergic neurons) in *Drosophila* by triggering oxidative stress [64–67]. Thus, xenobiotics are important to model PD in the fly. As a result, the reconstruction of the KEGG-MetaCyc network supported an extended metabolic information about *D. melanogaster*. This may be promising in future analyses to shed light on the molecular mechanisms underlying a variety of human diseases.

We subsequently compartmentalized the KEGG-MetaCyc-specific metabolic network since it did not include compartment information. The orthology-based draft model contains eight intracellular compartments (cytosol, nucleus, golgi apparatus, endoplasmic reticulum, mitochondria, mitochondrial intermembrane space, lysosome, and peroxisome) along with their Gene Orthology IDs. Considering these IDs, we first identified the compartments of *Drosophila* genes in the KEGG-MetaCyc network via available resources (COMPARTMENTS, FlyBase, GLAD, QuickGO, AmiGO 2, UniProt, Reactome, and a mass-spectrometry-based study). The subcellular localization information was mapped to each gene in the network. Missing compartments were predicted through CELLO2GO web server for the cut-off of 10^-5^ and so we generated a gene-compartment pair list. Using this approach, multiple compartments were assigned to many genes (Supplementary Table 3). This compartment dictionary allowed the assignment of subcellular localization(s) to each reaction based on the GPR associations. In this step, if a gene has multiple compartments, we assumed that the related reaction should be repeated in the model for each compartment. Thus, new reactions and metabolites were added to the network according to the subcellular locations of the corresponding gene, if necessary. In addition, if the genes catalyzing a reaction have different compartments, multiple compartments were assigned to this reaction and the corresponding genes were distributed based on their localizations. A general framework of the network compartmentalization process is illustrated in Figure 2 for the sake of clarity. The compartmentalized KEGG-MetaCyc-specific network consists of 1,077 genes and 3,511 metabolites involved in 2,015 enzymatic reactions.

**Figure 2.**
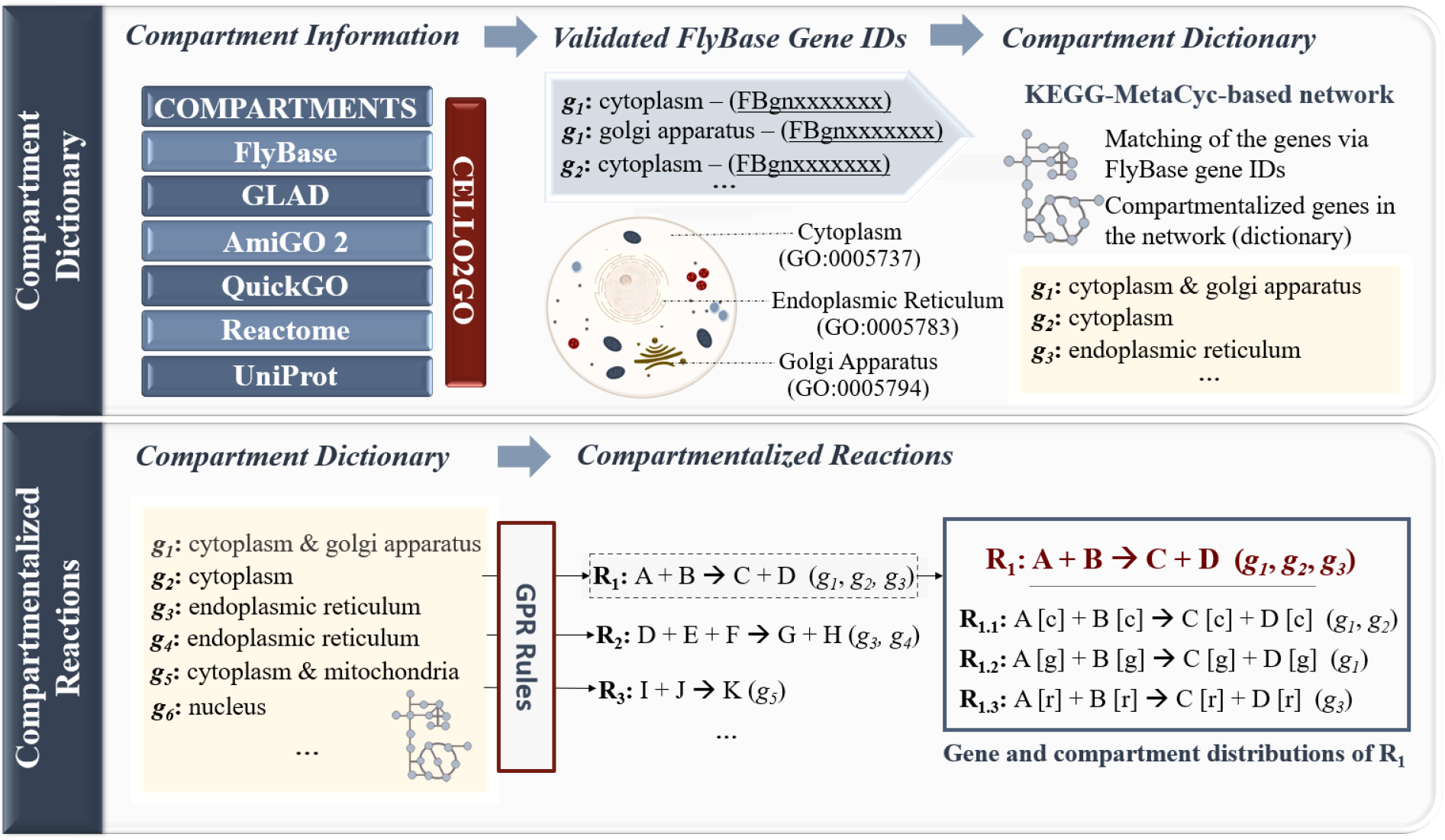
A general framework applied for the compartmentalization of KEGG-MetaCyc network. This process starts with the retrieval of compartment information along with validated FlyBase gene IDs from different sources. The compartment information is mapped to the genes in KEGG-MetaCyc network in order to generate a compartment dictionary. Based on the GPR associations, the compartment information is transferred from the genes to the reactions. In the given toy network, the reaction R_1_ provides the conversion of A and B metabolites to C and D metabolites, and it is catalyzed via the enzymes encoded by three different genes: *g_1_*, *g_2_*, and *g_3_*. In the compartmentalization process, R_1_ is added to the network in a repeated manner for each gene compartment (c: cytosol, g: golgi apparatus, and r: endoplasmic reticulum). The genes are subsequently distributed to the reactions (R_1.1_, R_1.2_, and R_1.3_) derived from R_1_ based on their compartment information.

### 3.2 Combining draft *Drosophila* metabolic networks and additional curations

The orthology-based draft *Drosophila* model was merged with the compartmentalized KEGG-MetaCyc network. Thus, the number of genes in the draft model was elevated to 2,398. Capacity of the merged model to produce biomass was assessed. We determined that many cofactors and vitamins could not be produced. A gap-filling algorithm was therefore applied to ensure the production of all biomass precursors. It allowed the addition of 29 new reactions (Supplementary Table 5) to the merged model. The newly added reactions enabled filling in the gaps related to cofactor and vitamin metabolism. In the next step, the gap-filled model was curated in terms of cholesterol and apocytochrome-C metabolism, GPR rules, and metabolic leaks as well as the common curation steps (Supplementary Note).

Cholesterol acts as the major structural component of *Drosophila* membrane and the precursor of steroid hormone. Steroid production is crucial in the regulation of developmental processes, which are required to generate an adult organism [68, 69] (Figure 3A). On the other hand, *D. melanogaster* has proved to be cholesterol auxotroph, that means inability for *de novo* cholesterol synthesis due to incomplete cholesterol biosynthesis pathway [70, 71]. In mammals, 3-hydroxy-3-methylglutaryl CoA reductase enzyme converts 3-hydroxy-3-methylglutaryl CoA into mevalonate. Using a set of enzymes, this compound is converted to farnesyl pyrophosphate. Santos and colleagues (2004) uncovered several fly orthologs catalyzing this pathway, which is branched to isoprenoid synthesis process (Figure 3B). The farnesyl pyrophosphate can be also directed to cholesterol synthesis branch in mammals. Most human genes involved in this branch (from farnesyl pyrophosphate to cholesterol) are not conserved in fly [47, 72] (Figure 3C). Hence, *Drosophila* must directly obtain sterols as dietary components [68, 70, 71]. Zhang and colleagues (2019) identified only two cholesterol synthesis genes in human (*SC4MOL* and *LBR*/*TM7SF2*) with *Drosophila* orthologs, whereas there are many non-orthologous human genes (*SQS*, *SQLE*, *LSS*, *CYP51A1*, *NSDHL*, *ERG27*, *DHCR24*, *EBP*, *SC5DL*, and *DHCR7*) [72] (Figure 3C).

**Figure 3.**
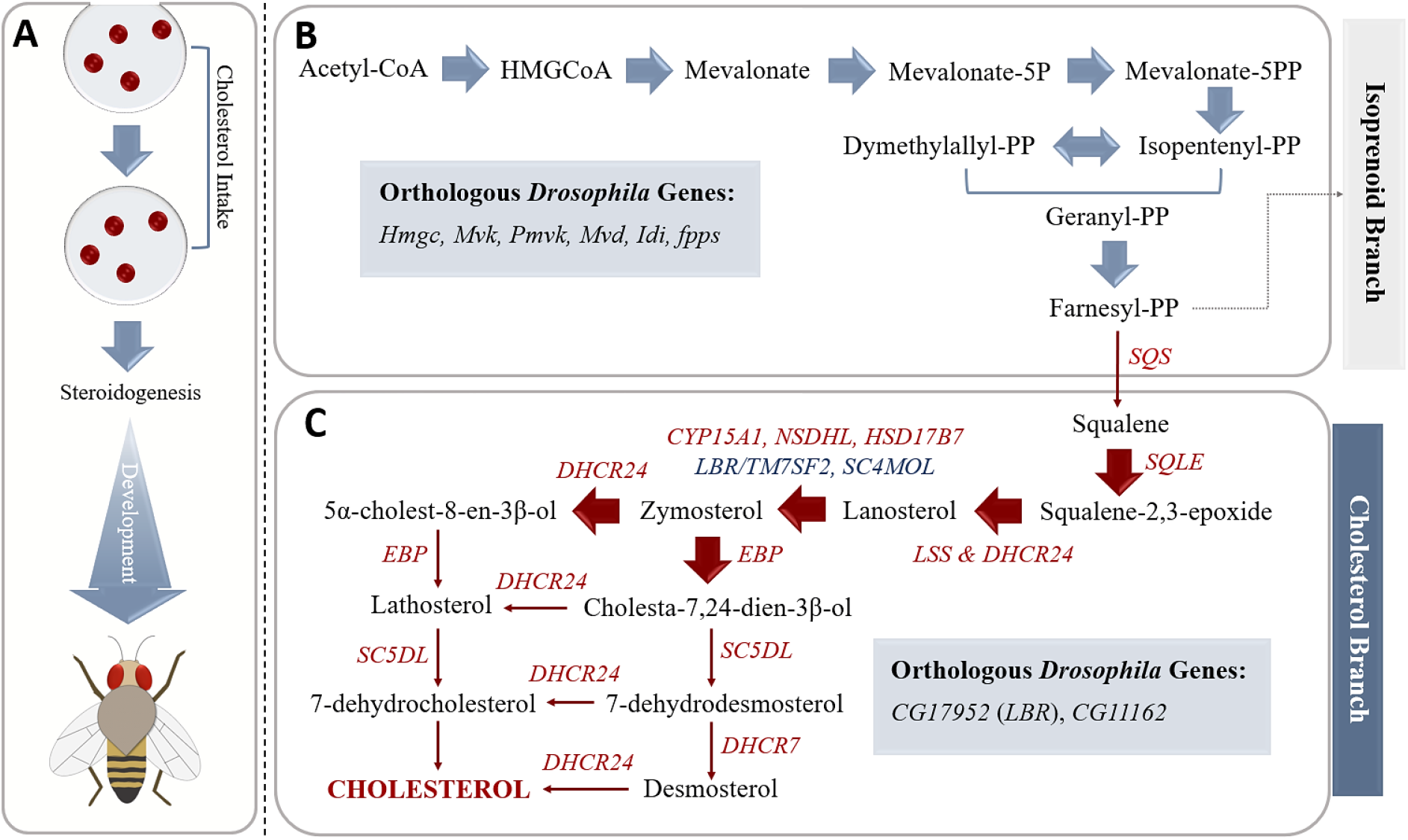
Cholesterol metabolism in *Drosophila melanogaster*. A) *Drosophila* needs cholesterol as a precursor to produce steroid hormones, which are crucial for the regulation of developmental processes. B) The initial steps of the cholesterol metabolic process including farnesyl-PP synthesis are conserved in all animals. The farnesyl-PP can be metabolized via two main pathways: isoprenoid branch and cholesterol branch in mammals. However, *Drosophila* does not include many genes in the cholesterol branch (indicated in red) and conserved genes in the cholesterol metabolism are indicated in blue. These metabolic gaps underlie cholesterol auxotrophy in the fruit fly.

We determined the cholesterol biosynthesis reactions in the draft *Drosophila* model that correspond to the inactive cholesterol branch. For these reactions, we first identified the corresponding *Drosophila* genes that were incorrectly defined as human orthologs. Accordingly, two *Drosophila* genes were found to be assigned as the orthologs of *NSDHL* gene encoding sterol-4-alpha-carboxylate 3-dehydrogenase (decarboxylating) and *DHCR7* gene encoding 7-dehydrocholesterol reductase in human. Of these *Drosophila* genes, *CG7724* (FBgn0036698) associated with steroid biosynthesis was matched with *NSDHL* (ENSG00000147383) at the maximum DIOPT score of 3. Since *Drosophila* was reported to lack *NSDHL* ortholog [72], we removed the corresponding reactions from the *Drosophila* model. These reactions are responsible for the synthesis of 3-keto-4-methylzymosterol, 5α-cholesta-8,24-dien-3-one, 4α-methyl-5α-cholesta-8-en-3-one, and 5α-cholesta-8-en-3-one compounds (Table 1). Another incorrectly assigned *Drosophila* gene, *LBR* (FBgn0034657) encoding lamin B receptor, was matched with *DHCR7* (ENSG00000172893) at the maximum DIOPT score of 5. Since *DHCR7* was also defined as the non-ortholog, we excluded the related reactions (HMR_1519 and HMR_1565) from the model (Table 1). HMR_1519 reaction is responsible for the formation of desmosterol from 7-dehydrodesmosterol and HMR_1565 allows the conversion of provitamin D3 (7-dehydrocholesterol) to cholesterol. Furthermore, eight non-conserved cholesterol biosynthesis reactions that were included into the *Drosophila* model in the gap-filling step were discarded from the model. These reactions are also listed in Table 1. In conclusion, we curated the cholesterol metabolism in the *Drosophila* model through the removal of the non-conserved human reactions. This step was necessary to mimic the cholesterol auxotrophy of fly. In addition to the cholesterol metabolism, we revised apocytochrome-C metabolism. To do so, the missing metabolite, apocytochrome-C, was added to the HMR_4762 reaction associated with porphyrin metabolism. Four reactions were added to the *Drosophila* model from HMR2 for the proper metabolism of apocytochrome-C (see Section 2.1).

**Table 1.**
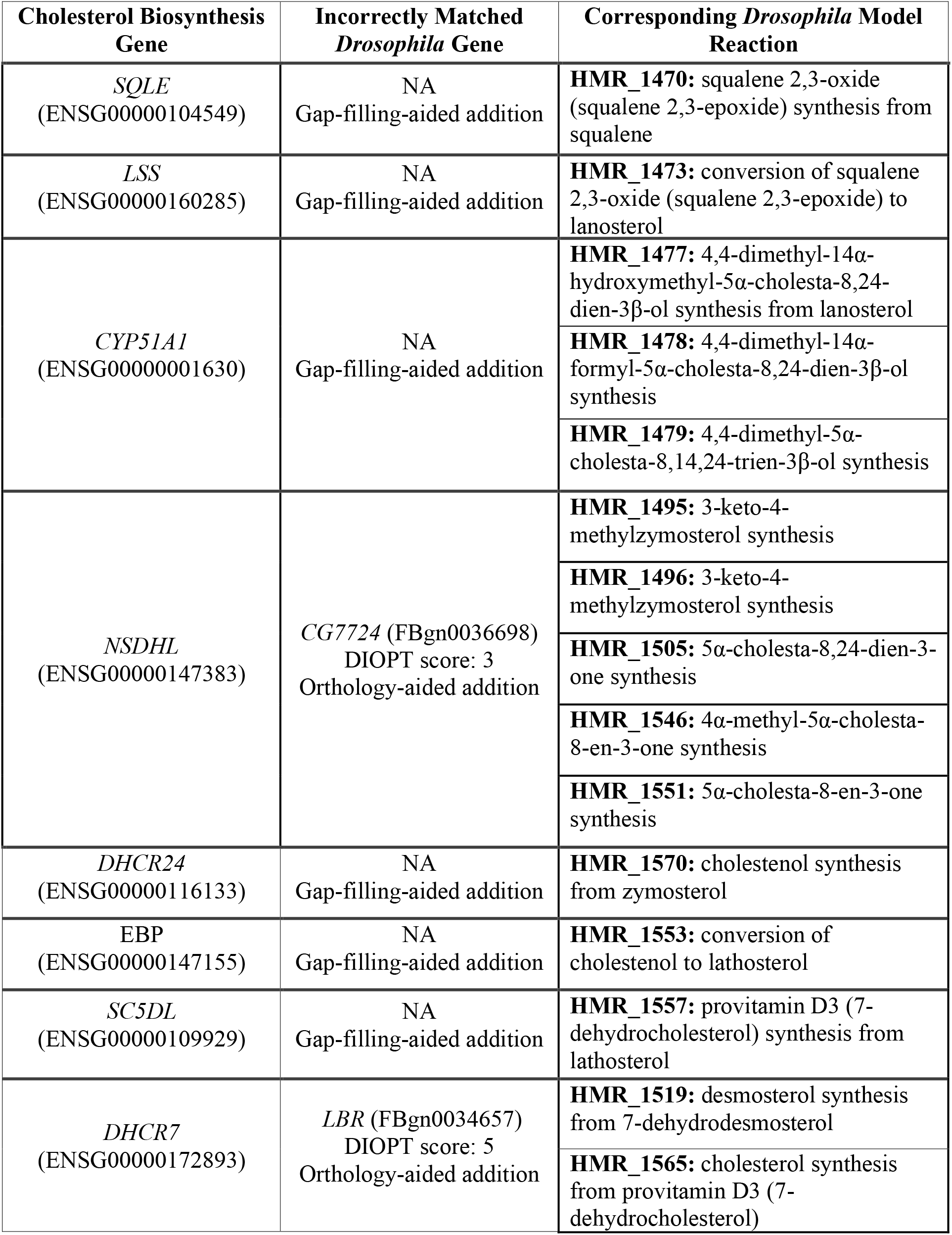
Metabolic network-driven investigation of non-conserved cholesterol biosynthesis reactions between human and fly. The non-conserved reactions that were incorrectly included in the draft *Drosophila* model via gap-filling or orthology-aided approach are determined based on the non-orthologous human genes in literature. These reactions are excluded from the draft *Drosophila* model in the next step.

Additionally, we curated the GPR rules lacking protein complex information. This was achieved based on the *Drosophila* network including 556 protein complexes, which was developed by Guruharsha and colleagues [35]. We linked the fly genes with ‘AND’ operators if they are found in the same protein complex. Based on FlyBase gene group list and previous studies [36, 37, 46–48, 38–45], we also curated especially energy metabolism-related gene associations such as mitochondrial complex I-V. The FlyBase gene groups consist of manually curated members within distinct gene families, the subunits of protein complexes, and other functional sets of the genes [39, 73]. We linked the genes encoding complex subunits (except for the paralogous genes) with ‘AND’ operators. This further refined the protein complex information in the GPR associations and facilitated the addition of the missing genes in the complexes. In particular, the modifications in mitochondrial complexes are crucial for the improved representation of the energy metabolism in condition-specific metabolic networks. Since impairments in mitochondrial function can lead to disrupted cellular phenomena (e.g., defective energy metabolism, elevated level of reactive oxygen species, and altered apoptotic signals), mitochondrial dysfunction was reported among the well-known causes of many neurodegenerative disorders [43, 74, 75]. Therefore, curation of the related GPR associations is important for more accurate characterization of such diseases. For accurate model simulations, we also updated biomass formation reaction based on FlySilico model [12] and literature (see Section 2.2).

The updated GMN model was further revised through the common curation steps explained in Supplementary Note. One significant modification is the removal of duplicated reactions. The duplicated reactions were identified via reaction comparisons by ignoring H^+^, H_2_O, and free inorganic phosphate (P_i_). Given that a relatively flexible compartmentation procedure was applied in KEGG-MetaCyc network, compartment information was excluded for the comparisons between KEGG-MetaCyc-derived reactions and template human model-derived reactions in the draft *Drosophila* model. Thus, we eliminated incorrectly compartmentalized KEGG-MetaCyc reactions from the *Drosophila* model by carefully examining each reaction match. This also enabled the curation of leaking ATP problem (i.e., spontaneous energy production even without any nutrient uptake) in the model via the removal of incorrectly compartmentalized uridine 5’-monophosphate phosphohydrolase, succinate dehydrogenase (ubiquinone), and NADH-dehydrogenase reactions. We further investigated the presence of other leaking energy metabolites in the draft model and identified four additional leaking metabolites including NADH, NADPH, FADH_2_, and H^+^. We subsequently performed multiple reaction deletion simulations via FBA to identify the reactions associated with metabolic leaks. These reactions were manually examined, and they were classified as incorrectly compartmentalized reactions or redundant reactions. The reason of this metabolic redundancy is the undetected duplicated reactions derived from KEGG-MetaCyc model and template human model due to small differences in metabolite names. Therefore, we removed these reactions from the model to block metabolite leakage. The final model called iDrosophila1 includes 8,230 reactions (5,787 enzymatic and 2,443 non-enzymatic), 6,990 metabolites, and 2,388 genes.

The iDrosophila1 reactions with missing subsystem (pathway) information were investigated through the KEGG database. If we could access pathway information of any reactions, these pathways were included in the model by denoting them as ‘KM pathways’. Analysis of pathways associated with iDrosophila1 reactions points to the dominance of major biological processes such as lipid, amino acid, and nucleotide metabolisms as well as cholesterol ester metabolism (Figure 4A). Especially lipid metabolic pathways were found to be prevalent. Cholesterol ester and triacylglycerol are the storage lipids accumulated in *Drosophila* fat body cells [76]. These dietary lipids are converted into free fatty acids, sterols, and monoacylglycerols that are absorbed by the cells of fly intestine under normal feeding conditions. Then, the resynthesized triacylglycerols are packaged into lipoproteins together with carrier proteins, cholesterols, and cholesterol esters for transportation along body. Thus, they can be used or stored by the tissues including adipose and liver. The presence of excess lipids induces the utilization of cholesterol esters and triacylglycerols as energy-supplying fuels [77]. Xenobiotic metabolism was determined as another prominent pathway in iDrosophila1. As highlighted before, xenobiotics are natural or synthetic life-threatening compounds that must be handled by animal cells via sequestration or metabolic degradation. Modification of the xenobiotics by phase I enzymes (e.g., cytochrome P450 monooxygenase and esterases) occurs in the first step of their metabolic detoxification [60, 61, 78]. In addition to the pathway information, we examined the compartment distribution in the model. We observed a similar compartment distribution profile between the template human model (Figure 4B) and iDrosophila1 (Figure 4C). Especially cytosolic reactions and transport reactions were found to have high frequencies in both models, and they were followed by mitochondrial distribution (Figure 4B and 4C).

**Figure 4.**
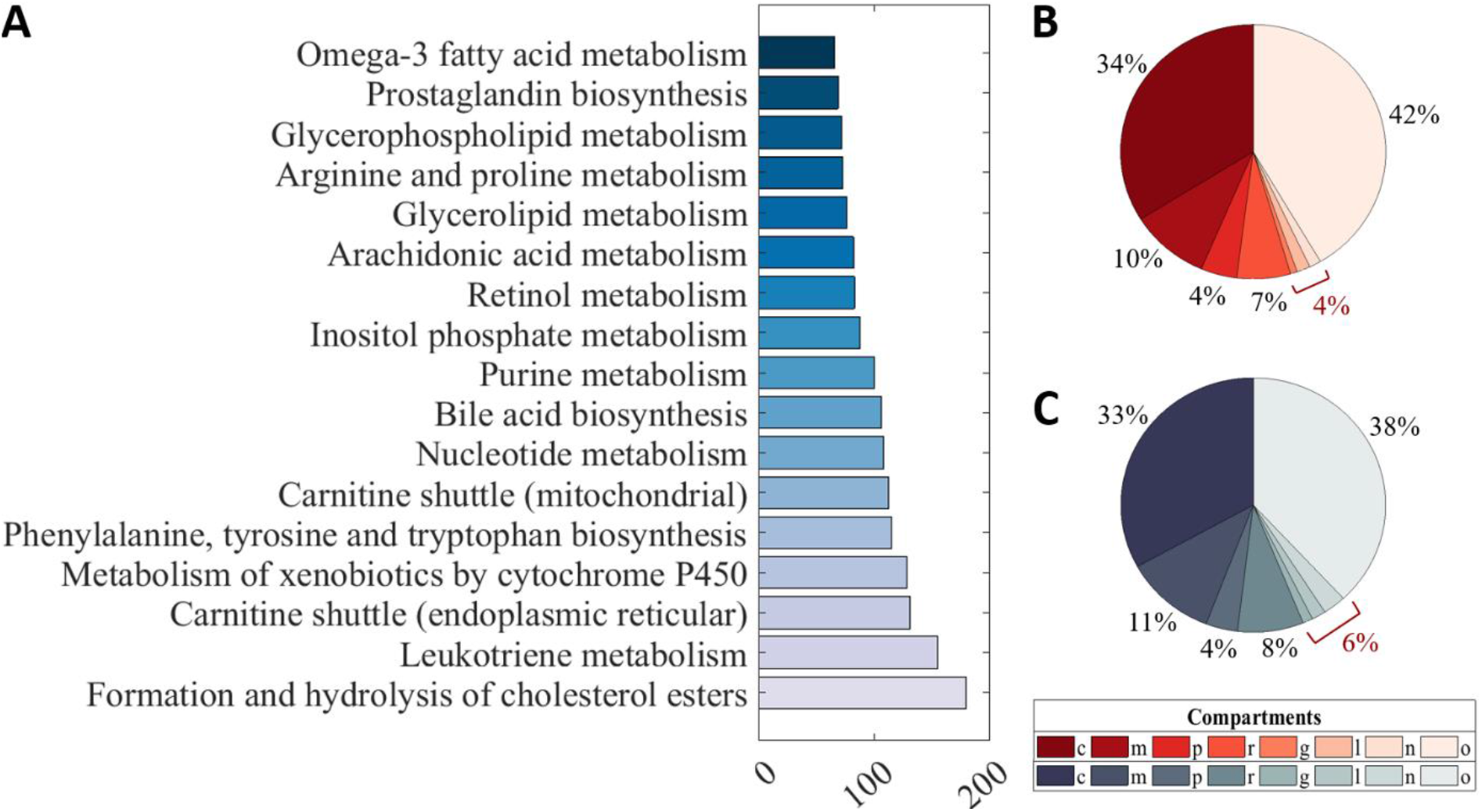
Distribution of the pathway and compartment information in iDrosophila1 model. A) Metabolic pathways in the model are ordered according to their frequency and top ten pathways are represented. Pie chart indicates the percentage of compartments in the B) template human model and C) iDrosophila1. The sum frequency values (given in red) are shown for the compartments with low frequencies. (Abbreviations: c, cytosol; m, mitochondria; p, peroxisome; r, endoplasmic reticulum; g, golgi apparatus; l, lysosome; n, nucleus; o, others (exchange and transport reactions)).

### 3.3 Prediction of growth phenotypes under increasing cholesterol and aspartate levels

In nature, *Drosophila* feeds on fermenting fruits containing high amounts of ethanol and organic acids. On the other hand, a simple diet of sucrose, lyophilized yeast, and weak organic acids was reported as sufficient to raise this organism in laboratory conditions. To mimic this dietary restriction, Piper and colleagues established the HD with essential and nonessential amino acids, vitamins, cholesterol, sucrose, several metal ions, and lipid precursors. This diet composition was reported to be convenient for the development of fruit flies [34]. Schönborn and colleagues simulated larval fly growth on the HD with a yeast-like amino acid ingredient. The researchers reported the predicted larval growth rate as 0.088 h^-1^ via FlySilico model by employing a set of constraints for the uptake rates [12]. Using iDrosophila1 model, we also analyzed the growth of *D. melanogaster* with the same uptake constraints (Supplementary Table 4). For the compatibility with FlySilico, we fixed the flux of non-growth associated maintenance reaction to the estimated value (8.55 mmol ATP g^-1^ h^-1^) [12]. We subsequently performed FBA simulation with growth maximization under the expanded HD condition. The growth rate was predicted to be 0.040 h^-1^ [12].

As aforementioned, cholesterol has crucial roles in membrane structure and signaling processes but flies are unable to synthesize this essential compound [71]. To properly mimic the cholesterol auxotrophic phenotype of *Drosophila*, we introduced several modifications in iDrosophila1 (Table 1) relying on literature [47, 72]. The capability of the model to accurately predict this phenotypic property was subsequently assessed. To this end, we allowed flexible intake of the expanded HD compounds by limiting maximum sucrose uptake rate to ∼ 2.212 mmol g^−1^h^−1^ and supplying the remaining metabolites at a certain ratio (see Section 2.3). In this condition, optimal growth rate was predicted as ∼ 0.774 h^-1^. We subsequently analyzed the growth profile across varying cholesterol levels from zero to ∼1/10 of the sucrose uptake rate. As expected, we did not observe biomass formation in the absence of cholesterol in the diet. Increasing level of the cholesterol positively induced biomass formation until reaching optimal growth rate (Figure 5A). We also performed this simulation through Fruitfly1 and FlySilico models by applying the same constraints to limit the consumption of the HD compounds. Fruitfly1 failed to grow under expanded HD condition. Therefore, three additional substances (lipoic acid, linoleate, and linolenate) were supplied to allow biomass formation. In addition, the right-hand side of biomass formation reaction in Fruitfly1 (version 1.1.0) is missing ADP that is required to establish balance between ATP and ADP. Therefore, we added this compound to the biomass formation reaction of this model for all simulations covered in this study. Note that no biomass production was obtained via Fruitfly1 while the measured uptake flux boundaries were introduced [12] (Supplementary Table 4). On the other hand, use of the flexible boundary constraints (see Section 2.3) triggered the growth rate of 0.450 h^-1^. FlySilico simulation, on the other hand, resulted in the maximum growth rate of 0.057 h^-1^ for the flexible uptake rates. In the next step, we examined the growth profiles of these models across the increasing cholesterol levels. Increasing cholesterol intake triggered elevated growth rate in FlySilico (Figure 5B) in agreement with the iDrosophila1 simulation while this phenotypic feature could not be simulated by Fruitfly1 (Figure 5C).

**Figure 5.**
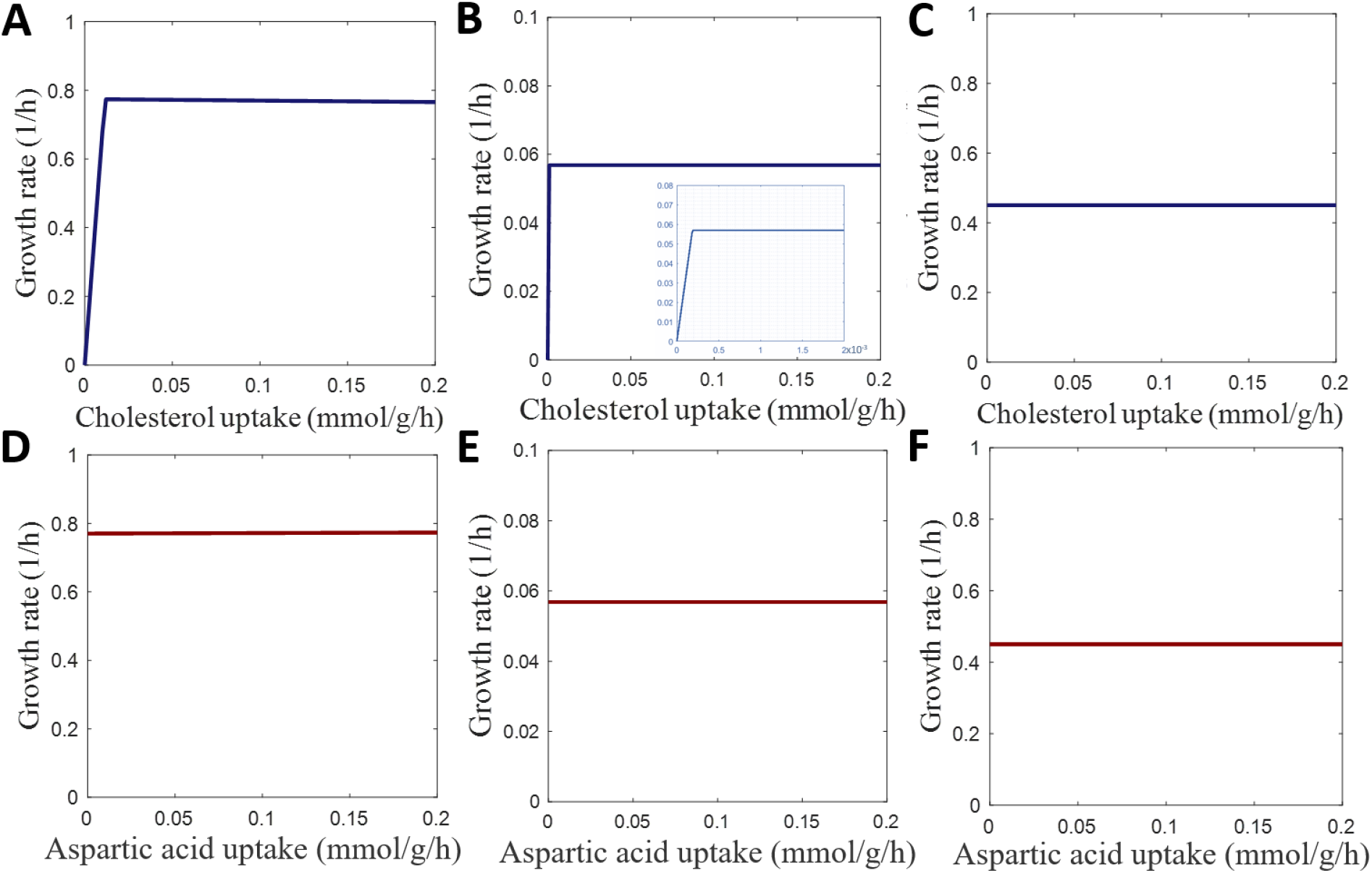
Growth profiles of the *Drosophila* under expanded HD condition supplied with elevating cholesterol and aspartate levels. A) iDrosophila1, B) FlySilico and C) Fruitfly1 models are used to characterize cholesterol-dependent changes in the proliferation. B) The zoom-out view associated with FlySilico simulations demonstrates a smaller uptake range of the cholesterol to clearly represent the relationship between exogenous cholesterol supplementation and growth profile. D), E) and F) display the effect of aspartate levels on the growth rates in the simulations by means of iDrosophila1, FlySilico, and Fruitfly1 models, respectively.

We also investigated the effect of varying aspartate levels on the fly growth under expanded HD condition. *Drosophila* contains ten essential amino acids, which were reported as commonly essential between mammals and insects, except for arginine [79, 80]. Aspartate is among the non-essential amino acids whose omission were reported to have no detrimental effect on the lifespan of *Drosophila* [34]. Using iDrosophila1 model, we revealed that biomass production was not affected from aspartate depletion thanks to its inherent aspartate biosynthesis system. Besides, supplementation with additional aspartate did not enhance the growth at a considerable level (Figure 5D). This is consistent with FlySilico (Figure 5E) and Fruitfly1 (Figure 5F) simulations (Figure 5E). Altogether, we confirmed the growth profile of fly across the varying levels of cholesterol and aspartate using iDrosophila1 model.

### 3.4 Prediction of essential *Drosophila* genes

Gene essentiality refers to the indispensability of genes for survival under specific growth conditions. We assessed the prediction performance of iDrosophila1 model by *in silico* single-gene knockouts. To do so, we blocked the corresponding reactions for the deletion of each gene. Essential genes were determined considering predicted growth rates. Accordingly, the genes whose deletions led to a significant reduction in growth rate were accepted as essential. For iDrosophila1 model, we elucidated essential and non-essential gene sets through FBA approach for unlimited intake of the expanded HD components, leading to 128 essential genes (Supplementary Table 6A) and performed GO and pathway enrichment analyses to provide an insight into these genes in terms of corresponding biological processes (Supplementary Table 6B) and pathways (Supplementary Table 6C). Unsurprisingly, these genes were found to be predominantly associated with biosynthetic processes of crucial cellular substances such as nucleotides, aminoacyl-tRNAs, amino acids, cofactors, and lipids. Enriched KEGG pathways were also identified to be consistent with these biological processes.

We further evaluated the iDrosophila1 performance through the comparison of the essentiality predictions with those obtained by other curated generic fly models. In this process, we introduced two main modifications on Fruitfly1 model prior to gene essentiality analysis. First modification is related to RNA metabolism. Two diverse cytosolic RNA synthesis reactions are present in both iDrosophila1 (HMR_7161 and HMR_7162) and Fruitfly1 (MAR07161 and MAR07162) models. They use nucleoside triphosphates (NTPs) and nucleoside diphosphates (NDPs) as the substrates, respectively. Since both molecules contain phosphoanhydride bonds, they are energy sources to drive biochemical reactions. NTPs (ATP, GTP, UTP, and CTP) also serve as substrates for nucleic acid biosynthesis by DNA-directed RNA polymerase (RNAP) enzymes [81]. Each RNAP enables the synthesis of distinct RNA classes from the ribosomal RNAs to non-coding RNAs [82]. Fruitfly1 and iDrosophila1 have nuclear and mitochondrial genes encoding RNAP subunits. Another reaction related to the RNA metabolism is catalysed by polynucleotide phosphorylase (PNPase) in the presence of NDPs [81, 83]. These conserved enzymes are responsible for RNA turnover primarily through the degradation of mitochondrial RNA [83, 84]. Recently, ATP-dependent RNA helicase SUV3 and PNPase enzymes were proposed to form a minimal mitochondrial RNA degradosome complex for mRNA decay in *Drosophila* [83]. PNPases (EC 2.7.7.8) were demonstrated to participate in RNA polymerization (non–template-encoded RNA synthesis) leading the yield of high P_i_ level, which can also allow RNA phosphorolysis [81, 85]. On the contrary, NTP polymerization by RNAPs occurs irreversibly. In Fruitfly1 model, RNAP genes were found in the GPR rules of both NDP polymerization (MAR07162) and NTP polymerization (MAR07161) reactions, whereas only PNPase-encoding *Drosophila* gene (FBgn0039846/ CG11337) is responsible for the catalysis of NDP polymerization (HMR_7162) in iDrosophila1 model. To support this, Pajak and colleagues reported that CG11337 gene is the fly ortholog of human PNPase [83]. The PNPase-mediated NDP polymerization is also consistent with our template human model [18] and iHsa model [17]. Taken together, we updated the GPR rule of Fruitfly1 reaction (MAR07162) before *in silico* gene deletions by replacing the RNAPs with the PNPase. In addition, we modified the reversibility of this reaction to represent its reversible characteristics. The reaction reversibility was also curated in iDrosophila1 model.

The second modification is related to the GPR association of the nucleo-cytoplasmic DNA transport reaction in Fruitfly1 (MAR08639), where nuclear pore complex genes were assigned. Nuclear pore complexes comprise several copies of ∼30 different proteins known as nucleoporins that mediate macromolecular trafficking (e.g., transport of RNAs, proteins and ribosomal subunits) between nucleus and cytoplasm as well as free diffusion of water, sugars, and ions [86–88]. Despite the presence of multiple *Drosophila* nucleoporins (e.g., Nup98 and Nup62) showing dynamic chromatin binding behavior in nucleoplasm, they only mediate transcriptional regulations [88]. The artificial DNA transport reactions in iDrosophila1 and Fruitfly1 models aim to reflect the contribution of DNA to cytosolic biomass formation. Hence, we removed the nucleoporin genes assigned to the artificial nucleo-cytoplasmic DNA transport reaction in Fruitfly1. Similar to the template human model [18] and iHsa model [17], iDrosophila1 model does not contain any genes dedicated to the corresponding transport reaction (HMR_8639).

Based on the updated GPR rules, we also determined essential (Supplementary Table 7A and 7B) and non-essential gene sets under HD condition for Fruitfly1 and FlySilico. Of the predicted 128 essential genes, iDrosophila1-specific results (n = 64) were predominantly found to be associated with the metabolism of nucleotides, cofactors, lipids, amino acids, and aminoacyl-tRNAs. For Fruitfly1-specific results (n = 33), metabolic processes associated with vitamins, cofactors, amino acids, aminoacyl-tRNAs, and nucleotide sugars were shown to be significantly enriched. On the other hand, FlySilico model predicted only six essential genes involved in carbohydrate and amino acid metabolism. The essential gene sets predicted by each generic fly model were also compared with the gene essentiality dataset retrieved from OGEE database [55]. We quantified the ratio of correct and incorrect essentiality predictions relying on four confusion matrix categories (true positives (TP), false positives (FN), true negatives (TN), and false negatives (FN)). And, we estimated several predictive scores including sensitivity, specificity, accuracy, precision, F1 score, and MCC. Although accuracy and F1 score are among the most widely favorable adopted metrics, they can be misleading in the evaluation of binary classification for particularly imbalanced datasets (e.g., many TNs but few TPs, or vice versa) [90]. Yet, MCC accounts for good prediction results in all confusion matrix categories suggesting the use of this robust metric for imbalanced datasets, as well [90, 91]. Here, we compared the predictive accuracy of iDrosophila1, Fruitfly1, and FlySilico models considering the metrics listed in Table 2.

**Table 2.**
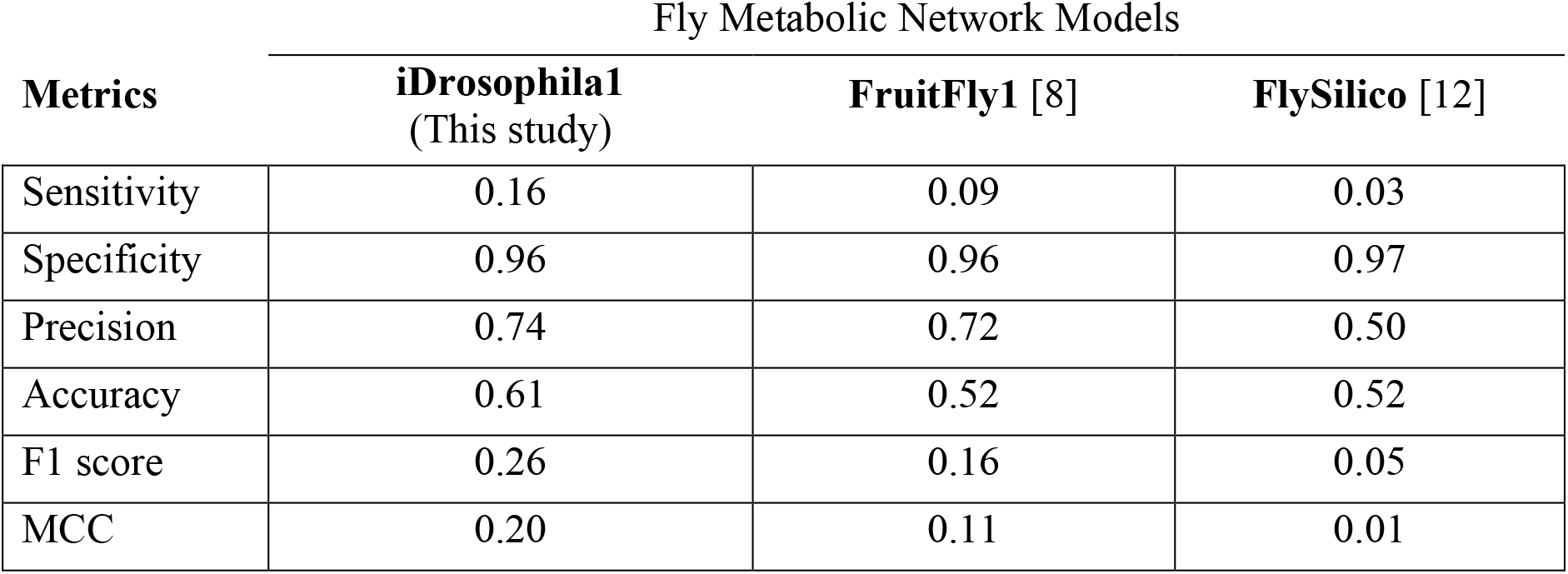
Comparison of the predictive metrics calculated for the generic fly models to assess their performances in terms of gene essentiality prediction.

| Metrics | Fly Metabolic Network Models |  |  |
| --- | --- | --- | --- |
|  | iDrosophila1<br>(This study) | FruitFly1 [8] | FlySilico [12] |
| Sensitivity | 0.16 | 0.09 | 0.03 |
| Specificity | 0.96 | 0.96 | 0.97 |
| Precision | 0.74 | 0.72 | 0.50 |
| Accuracy | 0.61 | 0.52 | 0.52 |
| F1 score | 0.26 | 0.16 | 0.05 |
| MCC | 0.20 | 0.11 | 0.01 |

The sensitivity of iDrosophila1 predictions were found to be considerably higher than the other fly models. Unsurprisingly, we revealed that FlySilico model had the worst sensitivity score due to the lack of protein complex information. In contrast to the low sensitivity values, the specificity results demonstrated that the fly models could correctly classify up to ∼96-97% of the non-essential genes. Based on precision values, iDrosophila1 (TP: 90 and FP: 32) and Fruitfly1 (TP: 64 and FP: 25) models were shown to predict the higher ratio of correct essential genes within the predicted essential gene sets in comparison with FlySilico (TP: 3 and FP: 3). Note that six essential genes predicted by iDrosophila1 could not be included in any confusion matrix categories due the lack of evidence while ten unclassified essential genes were identified via Fruitfly1 simulation. Since the binary metrics only consider two categories, we used additional metrics based on at least three confusion matrix categories. iDrosophila1 model was demonstrated to exhibit superior performance than the other models for these adopted metrics (accuracy, F1 score, and MCC). We found extremely low F1 score for FlySilico predictions. It was shown to be the highest for iDrosophila1 predictions, meaning the best compromise between sensitivity and precision. One drawback of the F1 score is the tendency of its value to converge into smaller values for low sensitivity or precision. Another issue is the misclassification potential in evaluating predictions under imbalanced prevalence [92]. We identified the frequency of correct non-essential genes (TN) as predominant in the predicted essential/non-essential gene sets for all fly models. Therefore, we calculated MCC scores, which vary in the interval of −1 and +1 indicating that a larger score reflects a better classification [90]. iDrosophila1 demonstrated a better predictive ability than the other models according to this metric, as well. Collectively, iDrosophila1 model represented comparable or better prediction results for all metrics.

### 3.5 Comprehensive metabolic profiling of Parkinson’s disease using iDrosophila1

We further analyzed iDrosophila1 to evaluate omics data-based prediction capacity of the model. In this regard, we investigated differential pathways in PD, which is the second most common age-related neurodegenerative disorder worldwide [64, 93]. This multifactorial disease has been attributed to different pathological hallmarks causing serious motor symptoms and non-motor symptoms [64, 94, 95]. Among the PD-causing factors, mitochondrial dysfunction plays a pivotal role in the pathogenesis of this disorder. Mitochondrial fusion and fission allow the exchange of cellular respiratory proteins and the selective removal of damaged mitochondria. Thus, they are crucial for a healthy mitochondrial homeostasis and neuroprotection [96–99]. In the fission process, a reduction in mitochondrial membrane potential promotes the accumulation of PINK1 (PTEN-induced novel kinase 1) on outer mitochondrial membrane [100]. Autophosphorylation and activation of these proteins recruit Parkin (a ubiquitin E3 ligase) from cytosol to mitochondria. A combined action of the Parkin and PINK1 triggers mitophagy-mediated degradation of the damaged mitochondria via phosphorylated stable polyubiquitin signals [101]. Any defects in this mitochondrial quality control process cause a PD-like phenotype (e.g., decreased lifespan, selective loss of dopaminergic neurons, locomotor abnormalities, and loss of olfaction) in *Drosophila* [102, 103].

Here, we characterized the metabolic alterations induced by *pink1* and *parkin* mutations using transcriptomic datasets, which were derived from young and middle-aged *Drosophila* models of PD [56]. Considering the altered transcriptome profiles (aka significant fold changes) of *pink1* and *parkin* mutant flies, regulated iDrosophila1 reactions were determined via ΔFBA approach (Figure 6A). The genes involved in these reactions were characterized through the detection of enriched metabolic pathways (Figure 6B; also, see Supplementary Table 8A-D for all enriched terms). Fundamental pathways associated with amino acid, nucleotide, and lipid metabolism were found to be commonly enriched for both young and middle-aged *pink1* and *parkin* mutants. Among the amino acid metabolic pathways, glycine and serine metabolism is especially prominent due to its role in *de novo* nucleotide biosynthesis pathway in the *Drosophila* [104]. The *de novo* nucleotide biosynthesis is primary strategy to sufficiently provide necessary DNA precursors for proliferating cells under normal growth conditions [105, 106]. In this process, a variety of substrates (glycine, glutamine, and 10-formyl-tetrahydrofolate) derived from pentose phosphate pathway, serine/glycine pathway, and one-carbon metabolism are used. Hence, folate supplementation was found to exhibit a stimulatory effect in *de novo* nucleotide biosynthesis pathways [107]. Based on personalized community-level modeling, Rosario and colleagues demonstrated a significant decrease of the folate production in patients with PD, correlated with the reduced bacterial folate biosynthesis [108]. Folate-supplemented diet was shown to be useful to prevent the loss of dopaminergic neurons in *pink1* mutant flies [104]. In agreement with the protective role of the folate in the reduction of mitochondrial defects [56, 104], we identified ‘folate biosynthesis’ as among the over-represented KEGG pathways for both *pink1* and *parkin* mutants in different age groups. Unsurprisingly, ‘purine and pyrimidine metabolism’ were also observed as significantly enriched. Another pathway related to folate metabolism is ‘riboflavin metabolism’, which was over-represented in all mutant groups. The water-soluble vitamin, riboflavin, is required by methylenetetrahydrofolate reductase enzymes, so it is crucial to ensure a healthy folate cycle. It also regulates the energy metabolism by acting as the precursor of FMN and FAD [109]. Thus, riboflavin exerts multiple protective functions against neurodegeneration by reducing glutamate excitotoxicity, oxidative stress, mitochondrial dysfunction, and NF-κB-induced neuroinflammation. Consistently, Parkin is responsible for stable glutamatergic synapses as well as the regulation of mitochondrial dynamics. In this regard, *parkin* mutation was reported to trigger an increased susceptibility to glutamate neurotoxicity, and this pathology is linked to the onset neurodegeneration of PD [110]. This may be evidence of the connection between riboflavin metabolism and mitochondrial quality control process (Figure 6B and Supplementary Table 8).

**Figure 6.**
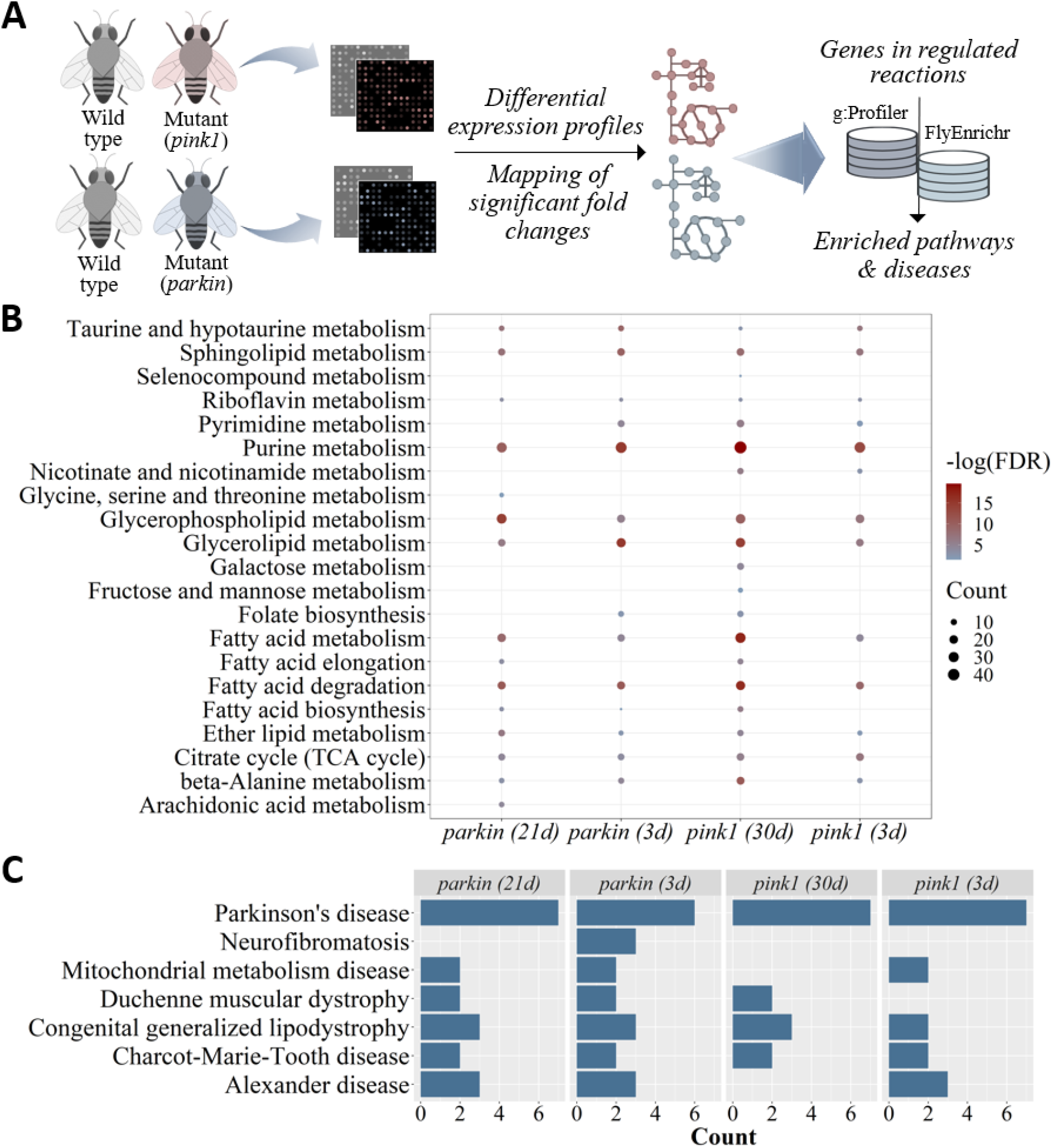
Identification of enriched pathways and diseases for the metabolic alterations induced by *pink1* and *parkin* mutations. **A)** Combinatory utilization of gene expression fold changes and metabolic network models are promising to explore differential reactions. For the genes involved in the regulated reactions, **B)** enriched pathways and **C)** diseases are illustrated for each mutant organism in different age group (3d: 3-day-old, 30d: 30-day-old, and 21d: 21-day-old). The enriched bubble chart shows the significant KEGG pathways (vertical axis) of the regulated genes discussed in the current study. The size of dots corresponds to the gene numbers in that pathway, and their colors represent enrichment significance. The darker red color indicates the higher significance. For the disease enrichment plot, vertical axis indicates the enriched diseases while the bars show the number of genes associated with the related disease.

Over-representation of ‘selenocompound metabolism’ was also determined for the 30-day-old *pink1* mutant (Figure 6B and Supplementary Table 8C). The essential micronutrient selenium plays a protective role against neurotoxicity by coping with oxidative stress and inflammation. It is capable of shaping gut microbiota, as well. An increased amount of *Bifidobacterium*, *Turicibacter*, and *Akkermansia* were reported in rodents via selenium supplementation [111]. Of these bacteria, *Akkermansia* was reported to be involved in gut barrier protection, immune modulation, and regulation of host metabolism [112]. On the other hand, selenium shows an age-dependent decline in human highlighting its possible relationship with a variety of diseases like PD and AD. Altogether, nutraceuticals and selenium-enriched functional foods (e.g., Baker’s yeast) have received a growing interest in recent years [111, 112]. Another over-represented metabolic process corresponding to neuroprotection is ‘β-alanine metabolism’ (Figure 6B and Supplementary Table 8). This non-proteinogenic amino acid is a precursor of carnosine dipeptide with diverse functions (e.g., proton buffering, metal chelation, antioxidation, regulation of calcium sensitivity, and muscle contractility), and its metabolic process was found to be enriched for all mutant groups. In PD patients, supplementation of carnosine dipeptide represents a therapeutic potential to improve the impaired motor activity [113]. Consistently, a relationship between the altered level of β-alanine and PD physiopathology was reported [114]. Although short term β-alanine treatment does not promote a substantial effect on the physical performance of PD patients [115], this compound may be promising due to its contribution to the enhanced levels of extracellular GABA and dopamine in substantia nigra [116]. In addition, taurine and β-alanine supplementation was shown to support muscle function in mice by increasing fatigue resistance in muscles and reducing contraction-induced oxidative stress [117]. Taurine is another β-amino acid involved in many physiological functions (e.g., neuromodulation, maintenance of calcium homeostasis, regulation of antioxidant and anti-inflammatory processes). At a high concentration in substantia nigra, it can particularly regulate dopamine release and the activity of dopaminergic neurons in healthy individuals. Che and colleagues demonstrated the taurine activity for the dopaminergic neuroprotection in the models of PD via inactivation of microglia-mediated neuroinflammation [118]. Here, ‘taurine and hypotaurine metabolism’ was found among the enriched pathways for both *pink1* and *parkin* mutants (Figure 6B and Supplementary Table 8). Thus, there is a further need to investigate the effect of long-term β-alanine and taurine supplementation on PD. More recently, nicotinamide riboside was also reported to have a potential neuroprotective effect against PD. NAD levels have been linked to ageing-related diseases and lifespan in the previous studies. Brakedal et al. showed that nicotinamide riboside regulates the levels of the genes associated with oxidative stress response, mitochondrial respiration, inflammatory response, histone acetylation, lysosomal processes, and proteasomal metabolism by elevating cerebral NAD levels in PD patients [119]. Consistently, we identified ‘nicotinate and nicotinamide metabolism’ as enriched in terms of differential fluxes in young and middle-aged *pink1* mutant flies (Figure 6B; also, see Supplementary Table 8A and 8C).

Lipid metabolic pathways including ‘fatty acid metabolism’, ‘fatty acid degradation’, ‘glycerolipid metabolism’, ‘glycerophospholipid metabolism’ ‘sphingolipid metabolism’, and ‘ether lipid metabolism’ were found to be enriched for *pink1* and *parkin* mutants in both age groups (Figure 6B and Supplementary Table 8). Brain includes a variety of lipids (e.g., fatty acids, triacylglycerols, sterols, and glycolipids) attributed to important functions in energy metabolism, signaling pathways, protein modifications, membrane activity and so on [120]. In recent reports, the lipid composition of synaptic vesicle membranes was defined as a key factor to regulate their interactions with α-synuclein (α-syn) proteins, whose aggregation is the main pathological hallmark of PD. These proteins preferentially interact with the vesicles including a certain lipid composition like arachidonoyl and docosahexaenoyl polyunsaturated fatty acids. Thus, lipid metabolism is prominent to modulate α-syn physiology and pathology [95, 120, 121]. Iljina and colleagues reported that arachidonic acid has a protective role against the generation of toxic beta-sheet structures causing the aberrant aggregation of α-syn proteins. This is because arachidonic acid supports the formation of alpha-helically-folded multimers of α-syn, and these structures exhibit higher resistance to fibril formation [122]. We identified the enrichment of arachidonic acid metabolism (Figure 6B). Furthermore, PINK1 and Parkin proteins accumulating at endoplasmic reticulum–mitochondria contact sites regulate inter-organelle communication associated with lipid metabolism, Ca^2+^ signaling, and mitophagy [123–125]. Overall, the mitochondrial activity is heavily linked to lipid metabolism, and altered lipidomic composition of the mitochondrial membranes in Parkin knock-out mice was documented by highlighting the tight relationship between lipid composition and mitochondrial activity [123]. In addition, fatty acid oxidation contributes acetyl-CoA production that is metabolized by tricarboxylic acid cycle. Tufi and colleagues demonstrated the downregulation of many tricarboxylic acid compounds in *pink1* mutant flies [104]. In a similar vein, we showed altered tricarboxylic acid cycle and carbon metabolism in *pink1* and *parkin* mutants (Figure 6B and Supplementary Table 8). In agreement with these results, *pink1* mutation was identified to be responsible for the reprogramming of glucose metabolism, which assists metabolic adaptation [126]. In particular, increased fructose levels were shown in PD patients by suggesting its potential role against oxidative stress through respiratory shifting in the early stages of PD. Carbon sources are also important to mediate protein modifications by glycation or glycosylation [127]. Here, we determined the enrichment of metabolic pathways linked with several glycating agents including fructose, galactose, and mannose in 30-day-old *pink1* mutant flies (Figure 6B and Supplementary Table 8B).

Lastly, we analyzed significantly enriched diseases associated with *pink1* and *parkin* mutations for each age group. Neurogenerative disorders, muscular, and mitochondrial diseases were found to be enriched by further confirming the accuracy of transcriptome-guided iDrosophila1 predictions (Figure 6C). Of the enriched disorders, ‘Duchenne muscular dystrophy’ is characterized by mitochondrial dysfunction and impaired mitophagy. Increased inflammation occurs due to the defective mitochondria, and it further triggers disease pathology corresponding to muscle damage and increased fibrosis in patients [128]. Similarly, Charcot-Marie-Tooth disease, which is a commonly inherited neurological disorder, leads to severe muscular deficits [129, 130]. It is associated with defective mitochondrial processes [129] in agreement with ‘mitochondrial metabolism disease’ and ‘Parkinson’s disease’ (Figure 6C). Together, these results also confirmed that we could successfully elucidate the PD-related differential *Drosophila* metabolism through the recently introduced ΔFBA approach. These preliminary findings are remarkable to indicate the capacity of iDrosophila1 in the discovery of context-specific pivotal metabolic alterations. Disease enrichment analyses further supported our model predictions by suggesting that this *Drosophila* model may be useful to gather insight into complicated human diseases.

## 4 Conclusion

In the current study, we reconstructed iDrosophila1 model using an orthology-based scoring approach. The orthology relationship between *Drosophila* and human genes, which are stored in FlyBase repository, was used to transfer metabolic information in the human model to the orthology-based fly model relying on high-confidence evidence scores. KEGG and MetaCyc databases enabled to expand metabolic coverage of the model. The most challenging task in combining metabolic knowledge from different sources is standardization issue. To overcome this problem, a variety of databases, tools, and literature were used. This step is crucial to avoid metabolic redundancy by supplying standardized metabolite and gene names. Each metabolic component of the model (reactions, metabolites, and genes) was further revised in the next curation steps. Furthermore, we used additional sources for the curation of metabolic information and protein complexes. Although such semi-automated standardization and curation steps are time-consuming, they are useful to improve organism-specific metabolic information in an accurate manner.

The predictive accuracy of iDrosophila1 was confirmed through phenotype-and gene essentiality-based analyses in a comparative manner with other curated generic fly models. In addition, gene expression fold change-mediated model analyses demonstrated the capacity of this model in the prediction of differential pathways over the course of Parkinson’s disease. Although *Drosophila* is a workhorse in experimental studies, there is still a gap in the use of this organism for comprehensive condition-specific metabolic modeling. Based on the increasing amounts of multi-omics data, the iDrosophila1 models contextualized by diverse disease-specific omics datasets may further contribute systems medicine. In this regard, such models may support novel insight into human disorders and biomarker detection. Moreover, community models have been developed to investigate multicellular metabolic interactions over the last decades. Available community models represent cell populations in single/multiple tissue(s), whole body, microbial communities, and host-microbial group interactions. Considering the dramatic impact of microbiota on human health and the low bacterial diversity of *Drosophila*, our metabolic network model can be also useful to elucidate metabolic interactions between *Drosophila* and gut microbiota. Overall, iDrosophila1 may provide avenues to advance fly metabolic modeling and better understand more complicated human metabolism.

## Author Contributions

MF performed orthology-based network reconstruction and metabolic network-based analyses. KP, TÇ conceived and designed the study. Manuscript was written by all authors.

## Acknowledgments

We would like to thank Katharina Zirngibl for providing the genome scale metabolic network of human, which was used as the template in the current study.

## Supplementary Notes

### Commonly applied curation steps for draft metabolic networks

Several common curation steps were introduced to increase the accuracy of each metabolic model, which was considered in the scope of this study. The networks were revised in terms of metabolic redundancy, stoichiometric consistency, name standardization, and missing/incomplete components. In this regard, each model component was carefully examined.

#### Reaction-centric curations

Reaction-centric curation ensured the absence of duplicated or stoichiometrically inconsistent reactions in the model. The duplicated reactions were identified in an iterative way by ignoring common currency metabolites (H^+^, H_2_O, and P_i_) from the model. When searching duplicated reactions derived from KEGG-MetaCyc-specific model and template human model [1], compartment information was also ignored due to the flexible compartmentalization process employed for KEGG-MetaCyc-specific model. This step was performed in a tightly coupled manner with the curation of gene-protein-reaction (GPR) rules for all models given in the current study. If the duplicated reactions are derived from different metabolic networks, the GPR rules of these reactions were manually examined and combined prior to the removal of one reaction by avoiding any gene loss. In addition, trivial or incomplete reactions with missing stoichiometric coefficients were removed from the models.

#### Gene-centric curations

Gene-centric curations allowed both standardization of gene names and the elimination of gene-based redundancy in models. Orthology-based draft *Drosophila* model was reconstructed using FlyBase gene IDs. On the other hand, KEGG-and MetaCyc-based models include FlyBase protein and annotation IDs. Therefore, all genes were denoted based on the current versions of FlyBase gene IDs for the compatibility between the models. However, this led to the emergence of redundant genes in GPR associations. Similarly, the conversion of Ensembl gene IDs into the FlyBase gene IDs in the orthology-based reconstruction of draft model caused the emergence of redundant genes due to the presence of multiple *Drosophila* orthologs for some human genes. To overcome this issue, the duplicated gene associations were curated. This step includes automatic detection of the duplicated gene associations (e.g., repeated isoenzymes and/or complex information) in each rule, updating this rule, followed by manual revision. Thus, each model was curated in such a way that it includes unique associations in each GPR rule.

#### Metabolite-centric curations

Metabolite-centric curations are based on the elimination of trivial and synonymous metabolites. The trivial metabolites, which were not associated with any reactions (no assigned stoichiometric coefficients), were investigated in all models and removed, if available. In addition, synonymous metabolites were identified based on two similar approaches to reduce metabolic redundancy. Metabolic redundancy can emerge due to the presence of duplicated (synonymous) metabolites within a single network and our first approach facilitated to cope with this issue. In addition, merging different models can lead to metabolic redundancy. To address this problem, we used the second approach.

In the first approach, synonymous metabolites within a model were interrogated based on compound names and IDs (KEGG [2], ChEBI [3], PubChem [4], and LIPID MAPS [5]). To this aim, compound information was collected via available metabolic networks and several web servers including Metabolite Translation Service [6], MBROLE 2.0 [7], MetaboAnalyst [8], and Chemical Translation Service [9] as well as available metabolic networks. Note that only three-star ChEBI IDs (parent IDs) manually curated by ChEBI team were kept in the compound list by excluding less reliable ChEBI IDs. Using this list, we subsequently generated a paired compound name and compound ID dictionary by mapping the related information to the metabolites in the given model. Based on these identifiers, we matched the model metabolites with each other. To increase the accuracy, the matched metabolites were considered as potentially synonymous only if they have the maximum number of common identifiers (Supplementary Figure 1). Based on available databases and literature, potential synonyms were subsequently revised via an extensive manual curation step to avoid wrong matches due to obsolete or incorrectly assigned IDs. For each metabolic match, one of the duplicated metabolites was removed from the model after the assembly of their stoichiometric coefficients in the model. This curation step was performed for the template human model, orthology-based draft *Drosophila* model, and KEGG-MetaCyc-specific network.

In the second approach, we uncovered synonymous metabolites found in different metabolic models to successfully merge these networks. This step was employed prior to combining orthology-based and KEGG-MetaCyc-specific metabolic networks to prevent any metabolic redundancy in the merged model. Similar to the previous approach, a compound name-ID pair dictionary was generated for each model (Supplementary Figure 1). Then, the metabolites in both models were mutually matched based on the specified identifiers. The same assumption was used to prioritize the matched metabolites. We selected the most reliable synonyms with the maximum number of common identifiers. After the manual confirmation of the potential synonyms, the names of synonymous metabolites in the KEGG-MetaCyc-specific network were replaced with the metabolite names in orthology-based draft *Drosophila* model. Thus, duplicated metabolites in the merged model could be easily distinguished and removed in the following step.

## Supplementary Figures

**Supplementary Figure 1.**
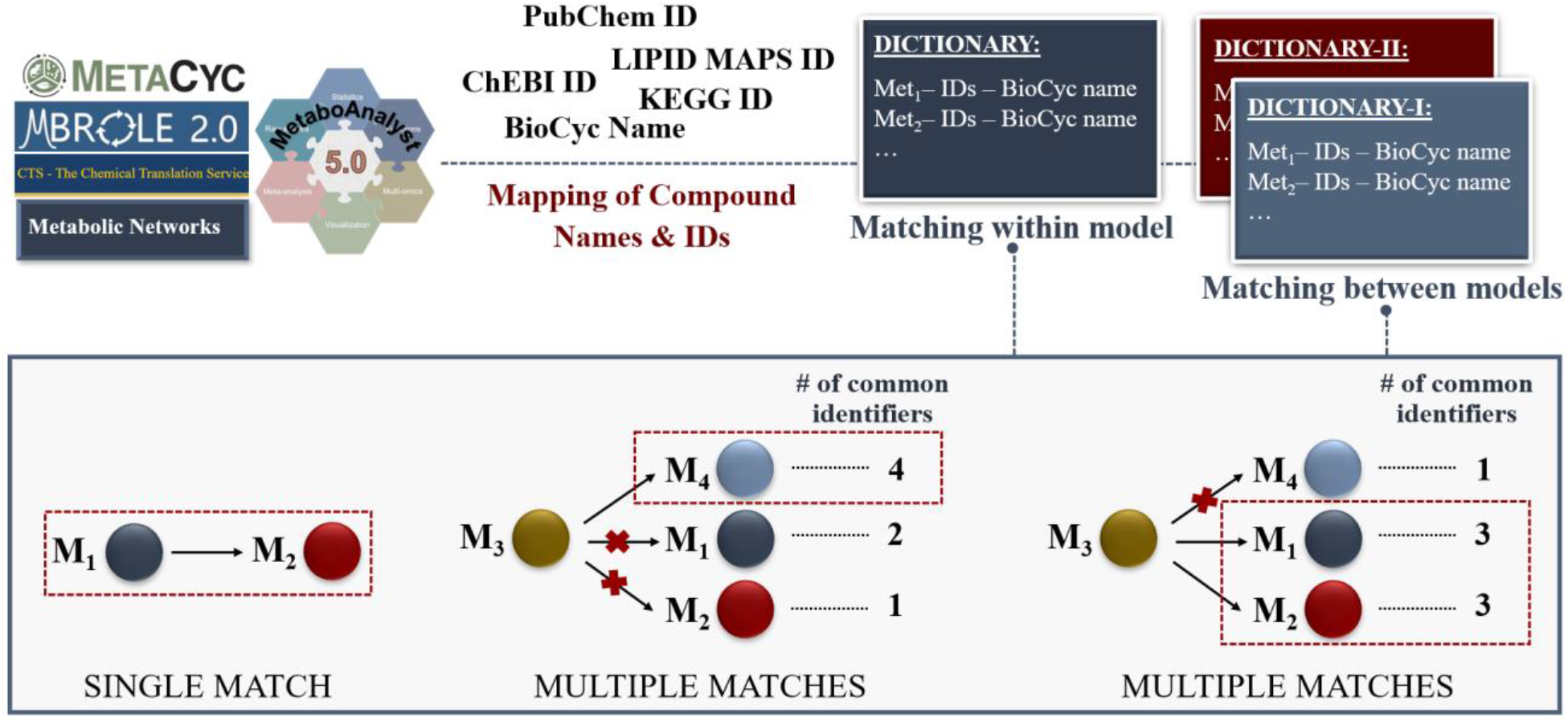
Identification and filtering of the synonymous metabolites through the generation of compound name and compound ID (KEGG, ChEBI, PubChem, and LIPID MAPS) pair dictionaries. To determine the synonymous metabolites within a model, a single dictionary is generated using several sources (Metabolite Translation Service, MBROLE 2.0, MetaboAnalyst, Chemical Translation Service, and available metabolic networks). Based on the matched identifiers in the dictionary, single and multiple hits can be determined. If a model compound has only one synonym as in the first illustrative example (see the metabolites M_1_ and M_2_), they should be accepted as synonymous metabolites. For the multiple matches, the number of the common identifiers is evaluated. If the metabolites M_3_ and M_4_ share the maximum number of the identifiers, they are accepted as synonyms. On the other hand, the multiple potential synonyms with the number of the maximum common identifiers are carefully examined to decide the correct compound pair. The same workflow steps are applied to identify the synonymous metabolites between different models. The only difference is to compare multiple dictionaries due to the creation of a different dictionary for each model.

## References

[1] Capo F., Wilson A., Di Cara F., (2019), “The intestine of Drosophila melanogaster: An emerging versatile model system to study intestinal epithelial homeostasis and host-microbial interactions in humans”, Microorganisms, 7(9).

[2] Mizuno H., Fujikake N., Wada K., Nagai Y., (2011), “α-Synuclein Transgenic Drosophila As a Model of Parkinson’s Disease and Related Synucleinopathies”, Park Dis, 2011(212706).

[3] Mirzoyan Z., Sollazzo M., Allocca M., Valenza A. M., (2019), “Drosophila melanogaster: A Model Organism to Study Cancer”, Front Genet, 10(51).

[4] Pandey U. B., Nichols C. D., (2011), “Human Disease Models in Drosophila melanogaster and the Role of the Fly in Therapeutic Drug Discovery”, Pharmacol Rev, 63(2), 411–436.

[5] Gu C., Kim G. B., Kim W. J., Kim H. U., Lee S. Y., (2019), “Current status and applications of genome-scale metabolic models”, Genome Biol, 20(121).

[6] Oberhardt M. A., Yizhak K., Ruppin E., (2013), “Metabolically re-modeling the drug pipeline”, Curr Opin Pharmacol, 13(5), 778–785.

[7] Nielsen J., Keasling J. D., (2016), “Engineering Cellular Metabolism”, Cell, 164(6), 1185–1197.

[8] Wang H., Robinson J. L., Kocabas P., Gustafsson J., Anton M., Cholley P.-E., Huang S., Gobom J., Svensson T., Uhlen M., Zetterberg H., Nielsen J., (2021), “Genome-scale metabolic network reconstruction of model animals as a platform for translational research”, Proc Natl Acad Sci, 118(30), e2102344118.

[9] Robinson J. L., Kocabaş P., Wang H., Cholley P., Cook D., Nilsson A., Anton M., Ferreira R., Domenzain I., Billa V., Limeta A., Hedin A., Gustafsson J., Kerkhoven E. J., Svensson L. T., Palsson B. O., Mardinoglu A., Hansson L., Uhlén M., Nielsen J., (2020), “An atlas of human metabolism”, Sci Signal, 13(624), eaaz1482.

[10] The Alliance of Genome Resources Consortium, (2020), “Alliance of Genome Resources Portal: unified model organism research platform”, Nucleic Acids Res, 48, 650–658.

[11] Büchel F., Rodriguez N., Swainston N., Wrzodek C., Czauderna T., Keller R., Mittag F., Schubert M., Glont M., Golebiewski M., Iersel M. Van, Keating S., Rall M., Wybrow M., Hermjakob H., Hucka M., Kell D. B., Müller W., Mendes P., Zell A., Chaouiya C., Saez-rodriguez J., Schreiber F., Laibe C., (2013), “Path2Models: large-scale generation of computational models from biochemical pathway maps”, BMC Syst Biol, 7(116).

[12] Schönborn J. W., Jehrke L., Mettler-altmann T., Beller M., (2019), “FlySilico: Flux balance modeling of Drosophila larval growth and resource allocation”, Sci Rep, 9(17156).

[13] Feala J. D., Coquin L., Mcculloch A. D., Paternostro G., (2007), “Flexibility in energy metabolism supports hypoxia tolerance in Drosophila flight muscle: metabolomic and computational systems analysis”, Mol Syst Biol, 3(99).

[14] Coquin L., Feala J. D., McCulloch A. D., Paternostro G., (2008), “Metabolomic and flux-balance analysis of age-related decline of hypoxia tolerance in Drosophila muscle tissue”, Mol Syst Biol, 4(233).

[15] Khodaee S., Asgari Y., Totonchi M., Karimi-jafari M. H., (2020), “iMM1865: A New Reconstruction of Mouse Genome-Scale Metabolic Model”, Sci Rep, 10(1), 6177.

[16] Sigurdsson M. I., Jamshidi N., Steingrimsson E., Thiele I., Palsson B. Ø., (2010), “A detailed genome-wide reconstruction of mouse metabolism based on human Recon 1”, BMC Syst Biol, 4(1), 140.

[17] Blais E. M., Rawls K. D., Dougherty B. V., Li Z. I., Kolling G. L., Ye P., Wallqvist A., Papin J. A., (2017), “Reconciled rat and human metabolic networks for comparative toxicogenomics and biomarker predictions”, Nat Commun, 8, 1–15.

[18] Zirngibl K., (2021), “Adaptability of Metabolic Networks in Evolution and Disease”. Ruperto Carola University.

[19] Larkin A., Marygold S. J., Antonazzo G., Attrill H., dos Santos G., Garapati P. V., Goodman J. L., Sian Gramates L., Millburn G., Strelets V. B., Tabone C. J., Thurmond J., (2021), “FlyBase: Updates to the Drosophila melanogaster knowledge base”, Nucleic Acids Res, 49(D1), D899–D907.

[20] Hu Y., Flockhart I., Vinayagam A., Bergwitz C., Berger B., Perrimon N., Mohr S. E., (2011), “An integrative approach to ortholog prediction for disease-focused and other functional studies”, BMC Bioinformatics, 12(357).

[21] Wang H., Marcis S., Hermansson D., Agren R., Nielsen J., Kerkhoven E. J., (2018), “RAVEN 2.0: A versatile toolbox for metabolic network reconstruction and a case study on Streptomyces coelicolor”, PLoS Comput Biol, 14(10), e1006541.

[22] Kanehisa M., Goto S., (2000), “KEGG: Kyoto Encyclopedia of Genes and Genomes”, Oxford Univ Press, 28(1), 27–30.

[23] Caspi R., Altman T., Dreher K., Fulcher C. A., Subhraveti P., Keseler I. M., Kothari A., Krummenacker M., Latendresse M., Mueller L. A., Ong Q., Paley S., Pujar A., Shearer A. G., Travers M., Weerasinghe D., Zhang P., Karp P. D., (2012), “The MetaCyc database of metabolic pathways and enzymes and the BioCyc collection of pathway/genome databases”, Nucleic Acids Res, 40, 742–753.

[24] Raudvere U., Kolberg L., Kuzmin I., Arak T., Adler P., Peterson H., Vilo J., (2019), “G:Profiler: A web server for functional enrichment analysis and conversions of gene lists (2019 update)”, Nucleic Acids Res, 47(W1), W191–W198.

[25] Binder J. X., Pletscher-frankild S., Tsafou K., Stolte C., Donoghue I. O., Schneider R., Jensen L. J., (2014), “COMPARTMENTS: unification and visualization of protein subcellular localization evidence”. Database (Oxford). doi: 10.1093/database/bau012.

[26] Hu Y., Comjean A., Perkins L. A., Perrimon N., Mohr S. E., (2015), “GLAD: an Online Database of Gene List Annotation for Drosophila”, J Genomics, 3, 75–81.

[27] Huntley R. P., Binns D., Dimmer E., Barrell D., Donovan C. O., Apweiler R., (2009), “QuickGO: a user tutorial for the web-based Gene Ontology browser”, Database (Oxford), 2009.

[28] Consortium T. G. O., (2015), “Gene Ontology Consortium: going forward”, Nucleic Acids Res, (Database issue), D1049–56.

[29] The Uniprot Consortium, (2021), “UniProt: the universal protein knowledgebase in 2021”, Nucleic Acids Res, 1–10.

[30] Fabregat A., Sidiropoulos K., Garapati P., Gillespie M., Hausmann K., Haw R., Jassal B., Jupe S., Korninger F., McKay S., Matthews L., May B., Milacic M., Rothfels K., Shamovsky V., Webber M., Weiser J., Williams M., Wu G., Stein L., Hermjakob H., D’Eustachio P., (2016), “The reactome pathway knowledgebase”, Nucleic Acids Res, 44(D1), D481–D487.

[31] Yu C.-S., Cheng C.-W., Su W.-C., Chang K.-C., Huang S.-W., Hwang J.-K., Lu C.-H., (2014), “CELLO2GO: A Web Server for Protein subCELlular LOcalization Prediction with Functional Gene Ontology Annotation”, PLoS One, 9(6), e99368.

[32] Tan D. J. L., Dvinge H., Christoforou A., Bertone P., Arias A. M., Lilley K. S., (2009), “Mapping Organelle Proteins and Protein Complexes in Drosophila melanogaster”, J Proteome Res, 8(6), 2667–2678.

[33] Heirendt L., Arreckx S., Pfau T., Mendoza S. N., Richelle A., Heinken A., Haraldsdóttir H. S., Wachowiak J., Keating S. M., Vlasov V., Magnusdóttir S., Ng C. Y., Preciat G., Alise Ž., Chan S. H. J., Aurich M. K., Clancy C. M., Modamio J., Sauls J. T., Noronha A., Bordbar A., Cousins B., Assal D. C. El, Valcarcel L. V, Apaolaza I., Ghaderi S., Ahookhosh M., Guebila M. Ben, Kostromins A., Sompairac N., Le H. M., Ma D., Sun Y., Wang L., Yurkovich J. T., Oliveira M. A. P., Vuong P. T., Assal L. P. El, Kuperstein I., Zinovyev A., Hinton H. S., Bryant W. A., Artacho F. J. A., Planes F. J., Stalidzans E., Maass A., Vempala S., Hucka M., Saunders M. A., Maranas C. D., Lewis N. E., Sauter T., Palsson B. Ø., Thiele I., Fleming R. M. T., (2019), “Creation and analysis of biochemical constraint-based models using the COBRA Toolbox v.3.0”, Nat Protoc, 14(3), 639–702.

[34] Piper M. D. W., Blanc E., Leitão-gonçalves R., Yang M., He X., Linford N. J., Hoddinott M. P., Hopfen C., Soultoukis G. A., Niemeyer C., Kerr F., Pletcher S. D., Ribeiro C., Partridge L., (2014), “A holidic medium for Drosophila melanogaster”, Nat Methods, 11(1), 100–105.

[35] Guruharsha K. G., Rual J. F., Zhai B., Mintseris J., Vaidya P., Vaidya N., Beekman C., Wong C., Rhee D. Y., Cenaj O., McKillip E., Shah S., Stapleton M., Wan K. H., Yu C., Parsa B., Carlson J. W., Chen X., Kapadia B., Vijayraghavan K., Gygi S. P., Celniker S. E., Obar R. A., Artavanis-Tsakonas S., (2011), “A protein complex network of Drosophila melanogaster”, Cell, 147(3), 690–703.

[36] Grant J., Saldanha J. W., Gould A. P., (2010), “A Drosophila model for primary coenzyme Q deficiency and dietary rescue in the developing nervous system”, DMM Dis Model Mech, 3(11–12), 799–806.

[37] Kemppainen K. K., Rinne J., Sriram A., Lakanmaa M., Zeb A., Tuomela T., Popplestone A., Singh S., Sanz A., Rustin P., Jacobs H. T., (2014), “Expression of alternative oxidase in Drosophila ameliorates diverse phenotypes due to cytochrome oxidase deficiency”, Hum Mol Genet, 23(8), 2078–2093.

[38] Tang X., Zhou B., (2013), “Iron Homeostasis in Insects: Insights from Drosophila Studies”, IUBMB Life, 65(10), 863–872.

[39] Attrill H., Falls K., Goodman J. L., Millburn G. H., Antonazzo G., Rey A. J., Marygold S. J., (2016), “FlyBase: establishing a Gene Group resource for Drosophila melanogaster”, Nucleic Acids Res, 44, 786–792.

[40] Allan A. K., Du J., Davies S. A., Dow J. A. T., (2005), “Genome-wide survey of V-ATPase genes in Drosophila reveals a conserved renal phenotype for lethal alleles”, Physiol Genomics, 22, 128–138.

[41] Kovacs L., Chao-Chu J., Schneider S., Gottardo M., Tzolovsky G., Dzhindzhev N. S., Riparbelli M. G., Callaini G., Glover D. M., (2018), “Gorab is a Golgi protein required for structure and duplication of Drosophila centrioles”, Nat Genet, 50(7), 1021–1031.

[42] Marygold S. J., Attrill H., Speretta E., Warner K., Magrane M., Berloco M., Cotterill S., McVey M., Rong Y., Yamaguchi M., (2020), “The DNA polymerases of Drosophila melanogaster”, Fly (Austin), 14(1–4), 49–61.

[43] Kay S., Anjaneyulu R., Naga M., Vidya S., Maximino G., (2020), “Insights from Drosophila on mitochondrial complex I”, Cell Mol Life Sci, 77(4), 607–618.

[44] Avval F. Z., Holmgren A., (2009), “Molecular Mechanisms of Thioredoxin and Glutaredoxin Ribonucleotide Reductase *”, J Biol Chem, 284(13), 8233–8240.

[45] Wahl M. C., Irmler A., Hecker B., Schirmer R. H., (2005), “Comparative Structural Analysis of Oxidized and Reduced Thioredoxin from Drosophila melanogaster”, J Mol Biol, 345(5), 1119–1130.

[46] Pavlovic Z., Bakovic M., (2013), “Regulation of Phosphatidylethanolamine Homeostasis—The Critical Role of CTP:Phosphoethanolamine Cytidylyltransferase (Pcyt2)”, Int J Mol Sci, 14(2), 2529–2550.

[47] Santos A. C., Lehmann R., (2004), “Isoprenoids Control Germ Cell Migration Downstream of HMGCoA Reductase”, Dev Cell, 6(2), 283–293.

[48] Van den Berghe G., Vincent M. F., Jaeken J., (1997), “Inborn errors of the purine nucleotide cycle: Adenylosuccinase deficiency”, J Inherit Metab Dis, 20(2), 193–202.

[49] Fritzemeier C. J., Hartleb D., Szappanos B., Papp B., Lercher M. J., (2017), “Erroneous energy-generating cycles in published genome scale metabolic networks: Identification and removal”, PLoS Comput Biol, 13(4), 1–14.

[50] Kubota-sakashita T. K. M., (2020), “Cardiolipin is essential for early embryonic viability and mitochondrial integrity of neurons in mammals”, FASEB J, 34(1), 1465–1480.

[51] Orth, Jeffrey D., Ines Thiele B. Ø. P., (2010), “What is flux balance analysis?”, Nat Biotechnol, 28(3), 245–248.

[52] Schellenberger J., Que R., Fleming R. M. T., Thiele I., Orth J. D., Feist A. M., Zielinski D. C., Bordbar A., Lewis N. E., Rahmanian S., Kang J., Hyduke D. R., Palsson B., (2011), “Quantitative prediction of cellular metabolism with constraint-based models: The COBRA Toolbox v2.0”, Nat Protoc, 6(9), 1290–1307.

[53] Pratapa A., Balachandran S., Raman K., (2015), “Fast-SL: An efficient algorithm to identify synthetic lethal sets in metabolic networks”, Bioinformatics, 31(20), 3299–3305.

[54] Mahadevan R., Schilling C. H., (2003), “The effects of alternate optimal solutions in constraint-based genome-scale metabolic models”, Metab Eng, 5(4), 264–276.

[55] Gurumayum S., Jiang P., Hao X., Campos T. L., Young N. D., Korhonen P. K., Gasser R. B., Bork P., Zhao X.-M., He L., Chen W.-H., (2020), “OGEE v3: Online Gene Essentiality database with increased coverage of organisms and human cell lines”, Nucleic Acids Res, 49(D1), D998–D1003.

[56] Celardo I., Lehmann S., Costa A. C., Loh S. H. Y., Martins L. M., (2017), “dATF4 regulation of mitochondrial folate-mediated one-carbon metabolism is neuroprotective”, Cell Death Differ, 4, 638–648.

[57] Law C. W., Chen Y., Shi W., Smyth G. K., (2014), “Voom: Precision weights unlock linear model analysis tools for RNA-seq read counts”, Genome Biol, 15(2), 1–17.

[58] Ravi S., Gunawan R., (2021), “ΔFBA-Predicting metabolic flux alterations using genome-scale metabolic models and differential transcriptomic data”, PLoS Comput Biol, 17(11), 1–18.

[59] Kuleshov M. V., Jones M. R., Rouillard A. D., Fernandez N. F., Duan Q., Wang Z., Koplev S., Jenkins S. L., Jagodnik K. M., Lachmann A., McDermott M. G., Monteiro C. D., Gundersen G. W., Ma’ayan A., (2016), “Enrichr: a comprehensive gene set enrichment analysis web server 2016 update”, Nucleic Acids Res, 44(W1), W90–W97.

[60] Misra J. R., Horner M. A., Lam G., Thummel C. S., (2011), “Transcriptional regulation of xenobiotic detoxification in Drosophila”, Genes Dev, 25(17), 1796–1806.

[61] Trinder M., Daisley B. A., Dube J. S., Reid G., (2017), “Drosophila melanogaster as a high-throughput model for host-microbiota interactions”, Front Microbiol, 8, 1–8.

[62] Chin-Chan M., Navarro-Yepes J., Quintanilla-Vega B., (2015), “Environmental pollutants as risk factors for neurodegenerative disorders: Alzheimer and Parkinson diseases”, Front Cell Neurosci, 9, 1–22.

[63] Bjørklund G., Dadar M., Chirumbolo S., Aaseth J., (2020), “The Role of Xenobiotics and Trace Metals in Parkinson’s Disease”, Mol Neurobiol, 57(3), 1405–1417.

[64] Nagoshi E., (2018), “Drosophila Models of Sporadic Parkinson’s Disease”, Int J Mol Sci, 19(11), 3343.

[65] Muñoz-Soriano V., Paricio N., (2011), “Drosophila Models of Parkinson’s Disease: Discovering Relevant Pathways and Novel Therapeutic Strategies”, Park Dis, 2011(520640).

[66] Coulom H., Birman S., (2004), “Chronic Exposure to Rotenone Models Sporadic Parkinson’s Disease in Drosophila melanogaster”, J Neurosci, 24(48), 10993–10998.

[67] Hosamani R., (2010), “Prophylactic treatment with Bacopa monnieri leaf powder mitigates paraquat-induced oxidative perturbations and lethality in Drosophila melanogaster”, Indian J Biochem Biophys, 47(2), 75–82.

[68] Danielsen E. T., Moeller M. E., Yamanaka N., King-jones K., Connor M. B. O., Rewitz K. F., (2016), “A Drosophila Genome-Wide Screen Identifies Regulators of Steroid Hormone Production and Developmental Timing”, Dev Cell, 37(6), 558–570.

[69] Niwa R., Niwa Y. S., (2011), “The fruit fly drosophila melanogaster as a model system to study cholesterol metabolism and homeostasis”, Cholesterol, 2011.

[70] Knittelfelder O., Prince E., Sales S., Fritzsche E., Wöhner T., Brankatschk M., Shevchenko A., (2020), “Sterols as dietary markers for Drosophila melanogaster”, Biochim Biophys Acta-Mol Cell Biol Lipids, 1865(7), 158683.

[71] Vinci G., Xia X., Veitia R. A., (2008), “Preservation of Genes Involved in Sterol Metabolism in Cholesterol Auxotrophs: Facts and Hypotheses”, PLoS One, 3(8), e2883.

[72] Zhang T., Yuan D., Xie J., Lei Y., Li J., Fang G., Tian L., Liu J., Cui Y., Zhang M., Xiao Y., Xu Y., Zhang J., Zhu M., Zhan S., Li S., (2019), “Evolution of the Cholesterol Biosynthesis Pathway in Animals”, Mol Biol Evol, 36(11), 2548–2556.

[73] Marygold S. J., Crosby M. A., Goodman J. L., Consortium T. F., (2016), “Using FlyBase, a Database of Drosophila Genes and Genomes”. In: Drosophila. pp 1–31.

[74] Compagnoni G. M., Di Fonzo A., Corti S., Comi G. P., Bresolin N., Masliah E., (2020), “The Role of Mitochondria in Neurodegenerative Diseases: the Lesson from Alzheimer’s Disease and Parkinson’s Disease”, Mol Neurobiol, 57, 2959–2980.

[75] Wu Y., Chen M., Jiang J., (2019), “Mitochondrial dysfunction in neurodegenerative diseases and drug targets via apoptotic signaling”, Mitochondrion, 49, 35–45.

[76] Liu Z., Huang X., (2013), “Lipid metabolism in Drosophila: Development and disease”, Acta Biochim Biophys Sin (Shanghai), 45(1), 44–50.

[77] Sieber M. H., Thummel C. S., (2012), “Coordination of triacylglycerol and cholesterol homeostasis by DHR96 and the drosophila lipa homolog magro”, Cell Metab, 15(1), 122–127.

[78] Amichot M., Tarès S., (2021), “The Foraging Gene, a New Environmental Adaptation Player Involved in Xenobiotic Detoxification”, Int J Env Res Public Heal, 14(18), 7508.

[79] Croset V., Schleyer M., Arguello J. R., Gerber B., Benton R., (2016), “A molecular and neuronal basis for amino acid sensing in the Drosophila larva”, Nat Publ Gr, 6(34871), 1–13.

[80] Manière G., Alves G., Berthelot-Grosjean M., Grosjean Y., (2020), “Growth regulation by amino acid transporters in Drosophila larvae”, Cell Mol Life Sci, 77(21), 4289–4297.

[81] Gottesman M. E., Mustaev A., (2019), “Ribonucleoside-5′-diphosphates (NDPs) support RNA polymerase transcription, suggesting NDPs may have been substrates for primordial nucleic acid biosynthesis”, J Biol Chem, 294(31), 11785–11792.

[82] Marygold S. J., Alic N., Gilmour D. S., Grewal S. S., (2020), “In silico identification of Drosophila melanogaster genes encoding RNA polymerase subunits”. MicroPubl Biol.

[83] Pajak A., Laine I., Clemente P., El-fissi N., Schober F. A., Maffezzini C., Javier C., Wibom R., Wredenberg A., (2019), “Defects of mitochondrial RNA turnover lead to the accumulation of double-stranded RNA in vivo”, PLoS Genet, 15(7), 1–25.

[84] Das S. K., Bhutia S. K., Sokhi U. K., Dash R., Azab B., Sarkar D., Fisher P. B., (2011), “Human polynucleotide phosphorylase (hPNPase old-35): an evolutionary conserved gene with an expanding repertoire of RNA degradation functions”, Oncogene, 30, 1733– 1743.

[85] Gasteiger E., Gattiker A., Hoogland C., Ivanyi I., Appel R. D., Bairoch A., Servet R. M., (2003), “ExPASy: the proteomics server for in-depth protein knowledge and analysis”, Nucleic Acids Res, 31(13), 3784–3788.

[86] Wente S. R., Rout M. P., (2010), “The Nuclear Pore Complex and Nuclear Transport”, Cold Spring Harb Perspect Biol, 2(10), 1–19.

[87] Ibarra A., Hetzer M. W., (2015), “Nuclear pore proteins and the control of genome functions”, Genes Dev, 29(4), 337–349.

[88] Kuhn T., Capelson M., (2018), “Nuclear Pore and Genome Organization and Gene Expression in Drosophila”. In: Nuclear Pore Complexes in Genome Organization, Function and Maintenance. Springer, Cham, pp 111–135.

[89] Munafò M., Lawless V. R., Passera A., Macmillan S., Bornelöv S., Haussmann I. U., Soller M., Hannon G. J., Czech B., (2021), “Channel nuclear pore complex subunits are required for transposon silencing in drosophila”, Elife, 10, 1–27.

[90] Chicco D., Jurman G., (2020), “The advantages of the Matthews correlation coefficient (MCC) over F1 score and accuracy in binary classification evaluation”, BMC Genomics, 21(6).

[91] Chicco D., Tötsch N., Jurman G., (2021), “The Matthews correlation coefficient (MCC) is more reliable than balanced accuracy, bookmaker informedness, and markedness in two-class confusion matrix evaluation”, BioData Min, 14(13), 1–22.

[92] Hand D. J., Christen P., Kirielle N., Christen P., (2021), “F*: an interpretable transformation of the F-measure”, Mach Learn, 110(3), 451–456.

[93] Aryal B., Lee Y., (2019), “Disease model organism for Parkinson disease: Drosophila melanogaster”, BMB Rep, 52(4), 250–258.

[94] Dung V. M., Thao D. T. P., (2018), “Parkinson’s Disease Model”, Adv Exp Med Biol, 1076, 41–61.

[95] Giguère N., Nanni S. B., Trudeau L., (2018), “On Cell Loss and Selective Vulnerability of Neuronal Populations in Parkinson’s Disease”, Front Neurol, 9(455).

[96] Sasaki S., Shirata A., Yamane K., Iwata M., (2004), “Parkin-positive autosomal recessive juvenile parkinsonism with α-synuclein-positive inclusions”, Neurology, 63(4), 678–683.

[97] Johansen K. K., Torp S. H., Farrer M. J., Gustavsson E. K., Aasly J. O., (2018), “Case of Parkinson’s Disease with No Lewy Body Pathology due to a Homozygous Exon Deletion in Parkin”, Case Rep Neurol Med, 2018(6838965), 19–21.

[98] Gouider-khouja N., Larnaout A., Amouri R., Sfar S., Belal S., (2003), “Autosomal recessive parkinsonism linked to parkin gene in a Tunisian family. Clinical, genetic and pathological study”, Park Relat Disord, 9(5), 247–251.

[99] Suda K., Mizuno Y., (1998), “Pathologic and biochemical studies of juvenile parkinsonism linked to chromosome 6q”, Neurology, 51(3), 890–2.

[100] Pathogenesis D., (2014), “Mitophagy Controlled by the PINK1-Parkin Pathway Is Associated with Parkinson’s”. In: Autophagy, Fourth Edi. Elsevier Inc., pp 227–238.

[101] Sekine S., Youle R. J., (2018), “PINK1 import regulation; a fine system to convey mitochondrial stress to the cytosol”, BMC Biol, 16(1), 2.

[102] Parker-Character J., Hager D. R., Call T. B., Pickup Z. S., Turnbull S. A., Marshman E. M., Korch S. B., Chaston J. M., Call G. B., (2021), “An altered microbiome in a Parkinson’s disease model Drosophila melanogaster has a negative effect on development”, Sci Rep, 11(1), 1–13.

[103] Xu Y., Xie M., Xue J., Xiang L., Li Y., Xiao J., Xiao G., Wang H. L., (2020), “EGCG ameliorates neuronal and behavioral defects by remodeling gut microbiota and TotM expression in Drosophila models of Parkinson’s disease”, FASEB J, 34(4), 5931–5950.

[104] Tufi R., Gandhi S., Castro I. P. De, Lehmann S., Angelova P. R., Dinsdale D., Deas E., Plun-favreau H., Nicotera P., Abramov A. Y., Willis A. E., Mallucci G. R., Loh S. H. Y., Martins L. M., (2014), “Enhancing nucleotide metabolism protects against mitochondrial dysfunction and neurodegeneration in a PINK1 model of Parkinson’s disease”, Nat Cell Biol, 16(2).

[105] Legent K., Mas M., Dutriaux A., Bertrandy S., Flagiello D., Delanoue R., Piskur J., Silber J., (2006), “In vivo analysis of Drosophila deoxyribonucleoside kinase function in cell cycle, cell survival and anti-cancer drugs resistance”, Cell Cycle, 5(7), 740–749.

[106] Holland C., Lipsett D. B., Clark D. V., (2011), “A link between impaired purine nucleotide synthesis and apoptosis in Drosophila melanogaster”, Genetics, 188(2), 359– 367.

[107] Villa E., Ali E. S., Sahu U., Ben-Sahra I., (2019), “Cancer cells tune the signaling pathways to empower de novo synthesis of nucleotides”, Cancers (Basel), 11(5), 1–20.

[108] Rosario D., Bidkhori G., Lee S., Bedarf J., Hildebrand F., Le Chatelier E., Uhlen M., Ehrlich S. D., Proctor G., Wüllner U., Mardinoglu A., Shoaie S., (2021), “Systematic analysis of gut microbiome reveals the role of bacterial folate and homocysteine metabolism in Parkinson’s disease”, Cell Rep, 34(9).

[109] Wong A. C. N., Dobson A. J., Douglas A. E., (2014), “Gut microbiota dictates the metabolic response of Drosophila to diet”, J Exp Biol, 217(11), 1894–1901.

[110] Marashly E. T., Bohlega S. A., (2017), “Riboflavin has neuroprotective potential: Focus on Parkinson’s disease and migraine”, Front Neurol, 8, 1–12.

[111] Callejón-Leblic B., Selma-Royo M., Collado M. C., Abril N., García-Barrera T., (2021), “Impact of Antibiotic-Induced Depletion of Gut Microbiota and Selenium Supplementation on Plasma Selenoproteome and Metal Homeostasis in a Mice Model”, J Agric Food Chem, 69(27), 7652–7662.

[112] Arias-Borrego A., Callejón-Leblic B., Calatayud M., Gómez-Ariza J. L., Collado M. C., García-Barrera T., (2019), “Insights into cancer and neurodegenerative diseases through selenoproteins and the connection with gut microbiota–current analytical methodologies”, Expert Rev Proteomics, 16(10), 805–814.

[113] Rezende N. S., Swinton P., de Oliveira L. F., da Silva R. P., da Eira Silva V., Nemezio K., Yamaguchi G., Artioli G. G., Gualano B., Saunders B., Dolan E., (2020), “The Muscle Carnosine Response to Beta-Alanine Supplementation: A Systematic Review With Bayesian Individual and Aggregate Data E-Max Model and Meta-Analysis”, Front Physiol, 11, 1–11.

[114] Solana-manrique C., Jos F., Torregrosa I., Palomino-schätzlein M., Hern C., Pineda-lucena A., Paricio N., (2022), “Metabolic Alterations in a Drosophila Model of Parkinson’s Disease Based on DJ-1 Deficiency”, Cells, 11(3), 1–17.

[115] Schön M., Mousa A., Berk M., Chia W. L., Ukropec J., Majid A., Ukropcová B., De Courten B., (2019), “The potential of carnosine in brain-related disorders: A comprehensive review of current evidence”, Nutrients, 11(6).

[116] Allman B. R., Biwer A., Maitland C. G., DiFabio B., Coughlin E., Smith-Ryan A. E., Ormsbee M. J., (2018), “The effect of short term beta alanine supplementation on physical performance and quality of life in parkinson’s disease: A pilot study”, J Exerc Physiol Online, 21(1), 1–13.

[117] Horvath D. M., Murphy R. M., Mollica J. P., Hayes A., Goodman C. A., (2016), “The effect of taurine and β-alanine supplementation on taurine transporter protein and fatigue resistance in skeletal muscle from mdx mice”, Amino Acids, 48(11), 2635–2645.

[118] Che Y., Hou L., Sun F., Zhang C., Liu X., Piao F., Zhang D., Li H., Wang Q., (2018), “Taurine protects dopaminergic neurons in a mouse Parkinson’s disease model through inhibition of microglial M1 polarization”, Cell Death Dis, 9(4).

[119] Brakedal B., Dölle C., Riemer F., Ma Y., Nido G. S., Skeie G. O., Craven A. R., Schwarzlmüller T., Brekke N., Diab J., Sverkeli L., Skjeie V., Varhaug K., Tysnes O. B., Peng S., Haugarvoll K., Ziegler M., Grüner R., Eidelberg D., Tzoulis C., (2022), “The NADPARK study: A randomized phase I trial of nicotinamide riboside supplementation in Parkinson’s disease”, Cell Metab, 34(3), 396–407.e6.

[120] Mori A., Imai Y., Hattori N., (2020), “Lipids: Key players that modulate α-synuclein toxicity and neurodegeneration in Parkinson’s disease”, Int J Mol Sci, 21(9).

[121] Chia S. J., Tan E., Chao Y., (2020), “Historical Perspective: Models of Parkinson’s Disease”, Int J Mol Sci, 21(7), 2464.

[122] Iljina M., Tosatto L., Choi M. L., Sang J. C., Ye Y., Hughes C. D., Bryant C. E., Gandhi S., Klenerman D., (2016), “Arachidonic acid mediates the formation of abundant alpha-helical multimers of alpha-synuclein”, Sci Rep, 6(33928), 1–14.

[123] Fais M., Dore A., Galioto M., Galleri G., Crosio C., Iaccarino C., (2021), “Parkinson’s disease-related genes and lipid alteration”, Int J Mol Sci, 22(14), 1–13.

[124] Barazzuol L., Giamogante F., Brini M., Calì T., (2020), “PINK1/ParkinMediatedMitophagy, Ca2+ Signalling, and ER–Mitochondria Contacts in Parkinson’s Disease”, Int J Mol Sci, 21(5), 1772.

[125] Gómez-Suaga P., Bravo-San Pedro J. M., González-Polo R. A., Fuentes J. M., Niso-Santano M., (2018), “ER-mitochondria signaling in Parkinson’s disease”, Cell Death Dis, 9(3).

[126] Requejo-Aguilar R., Lopez-Fabuel I., Fernandez E., Martins L. M., Almeida A., Bolaños J. P., (2014), “Pink1 deficiency sustains cell proliferation by reprogramming glucose metabolism through hif1”, Nat Commun, 5.

[127] Videira P. A. Q., Castro-Caldas M., (2018), “Linking glycation and glycosylation with inflammation and mitochondrial dysfunction in Parkinson’s disease”, Front Neurosci, 12, 1–20.

[128] Reid A. L., Alexander M. S., (2021), “The interplay of mitophagy and inflammation in duchenne muscular dystrophy”, Life, 11(7).

[129] Schiavon C. R., Shadel G. S., Manor U., (2021), “Impaired Mitochondrial Mobility in Charcot-Marie-Tooth Disease”, Front Cell Dev Biol, 9, 1–15.

[130] Nandini S., Conley Calderon J. L., Sabblah T. T., Love R., King L. E., King S. J., (2019), “Mice with an autosomal dominant Charcot-Marie-Tooth type 2O disease mutation in both dynein alleles display severe moto-sensory phenotypes”, Sci Rep, 9(1), 1–13.

## Supplementary References

[1] Zirngibl K., (2021), “Adaptability of Metabolic Networks in Evolution and Disease”. Ruperto Carola University.

[2] Kanehisa M., Goto S., (2000), “KEGG: Kyoto Encyclopedia of Genes and Genomes”, Oxford Univ Press, 28(1), 27–30.

[3] Degtyarenko K., De matos P., Ennis M., Hastings J., Zbinden M., Mcnaught A., Alcántara R., Darsow M., Guedj M., Ashburner M., (2008), “ChEBI: A database and ontology for chemical entities of biological interest”, Nucleic Acids Res, 36, 344–350.

[4] Kim S., Chen J., Cheng T., Gindulyte A., He J., He S., Li Q., Shoemaker B. A., Thiessen P. A., Yu B., Zaslavsky L., Zhang J., Bolton E. E., (2021), “PubChem in 2021: New data content and improved web interfaces”, Nucleic Acids Res, 49(D1), D1388–D1395.

[5] Fahy E., Sud M., Cotter D., Subramaniam S., (2007), “LIPID MAPS online tools for lipid research”, Nucleic Acids Res, 35, 606–612.

[6] Caspi R., Billington R., Ferrer L., Foerster H., Fulcher C. A., Keseler I. M., Kothari A., Krummenacker M., Latendresse M., Mueller A., Ong Q., Paley S., Subhraveti P., Weaver D. S., Karp D., (2016), “The MetaCyc database of metabolic pathways and enzymes and the BioCyc collection of pathway/genome databases”, Nucleic Acids Res, 44(1), 471–480.

[7] López-Ibáñez J., Pazos F., Chagoyen M., (2016), “MBROLE 2.0-functional enrichment of chemical compounds”, Nucleic Acids Res, 44(W1), W201–W204.

[8] Chong J., Soufan O., Li C., Caraus I., Li S., Bourque G., Wishart D. S., Xia J., (2018), “MetaboAnalyst 4.0: towards more transparent and integrative metabolomics analysis”, Nucleic Acids Res, 46, 486–494.

[9] Wohlgemuth G., Haldiya P. K., Willighagen E., Kind T., Fiehn O., (2010), “The Chemical Translation Service — a web-based tool to improve standardization of metabolomic reports”, Bioinformatics, 26(20), 2647–2648.

